# Temporal integration of narrative information in a hippocampal amnesic patient

**DOI:** 10.1101/713180

**Authors:** Xiaoye Zuo, Christopher J. Honey, Morgan D. Barense, Davide Crombie, Kenneth A. Norman, Uri Hasson, Janice Chen

**Affiliations:** Department of Psychological and Brain Sciences, Johns Hopkins University, Baltimore, MD 21218, USA; Department of Psychology, University of Toronto, Toronto, Ontario, M5S 3G3, Canada; Rotman Research Institute, Baycrest Hospital, Toronto, Ontario, M5S 3G3, Canada; Department of Biology II - Neurobiology, Ludwig Maximilian University of Munich, Großhaderner Str. 2, 82152 Planegg, Germany; Princeton Neuroscience Institute, Princeton University, Princeton, NJ 08544, USA; Department of Psychology, Princeton University, Princeton, NJ 08544, USA

## Abstract

Default network regions appear to integrate information over time windows of 30 seconds or more during narrative listening. Does this long-timescale capability require the hippocampus? Amnesic behavior suggests that the hippocampus may not be needed for online processing when input is continuous and semantically rich: amnesics can participate in conversations and tell stories spanning minutes, and when tested immediately on recently heard prose their performance is relatively preserved. We hypothesized that default network regions can integrate the semantically coherent information of a narrative across long time windows, even in the absence of the hippocampus. To test this prediction, we measured BOLD activity in the brain of a hippocampal amnesic patient (D. A.) and healthy control participants while they listened to a seven-minute narrative. The narrative was played either in its intact form, or as a paragraph-scrambled version, which has been previously shown to interfere with the long-range temporal dependencies in default network activity. In the intact story condition, D. A.’s moment-by-moment BOLD activity spatial patterns were similar to those of controls in low-level auditory cortex as well as in some high-level default network regions (including lateral and medial posterior parietal cortex). Moreover, as in controls, D. A.’s response patterns in medial and lateral posterior parietal cortex were disrupted when paragraphs of the story were presented in a shuffled order, suggesting that activity in these areas did depend on information from 30 seconds or more in the past. Together, these results suggest that some default network cortical areas can integrate information across long timescales, even in the absence of the hippocampus.

## Introduction

People with hippocampal damage are profoundly impaired in recalling information after a distraction (Milner, 1966), or “as soon as their attention shifts to a new topic” (Milner, 2005). At the same time, these amnesic individuals are able to engage in conversation, retain near-normal immediate recall for prose passages (Baddeley and Wilson, 2002), and tell globally coherent stories (Keven et al., 2018; Kurczek and Duff, 2011; Rosenbaum et al., 2009). How is this possible? One potential explanation is that, in the absence of major topic changes or surprises that create a distraction, amnesic individuals are able to rely on default network cortical regions for retention of semantically-rich information across time. Default network areas are proposed to carry slowly-changing information during continuous natural input such as stories and conversation (Hasson et al., 2015). In healthy people, these areas are functionally coupled to the hippocampus and may work together to accumulate, maintain, and integrate information across events. However, it is an open question whether the long-timescale capability of default network regions depends critically on interactions with the hippocampus (Chen et al., 2016).

In this study, we investigated whether default network areas can integrate information over tens of seconds even without the hippocampus by recording the brain activity of a hippocampal amnesic patient as he listened to an auditory story. Narratives that unfold over minutes are suitable to test such a question, as they require the integration of incoming information with relevant information presented many sentences ago. For example, the meaning of an incoming sentence, e.g., “X entered the bank”, will be different as a function of information presented earlier the story, e.g., X is a robber versus X is an investor. In previous studies, we demonstrated that (in neurotypical subjects) default network responses for a given paragraph of a story were changed if the preceding paragraph was changed by scrambling the order of paragraphs (Lerner et al., 2011); since each paragraph was on average 30 seconds long, the results indicate that default network activity patterns depend on information accumulated across at least the last 30 seconds of the story, i.e., long-timescale integration (**Supplementary Figure 1**). (Windows longer than 30 seconds have not been tested.) This scrambling technique was used with different window sizes to map a hierarchy of timescales across the cortex: default network areas had the longest timescales, intermediate areas along the superior temporal gyrus had sentence-length (hundreds of milliseconds) timescales, and early sensory areas were found to operate at timescales shorter than a word (tens of milliseconds) (Hasson et al., 2008; Honey et al., 2012; Lerner et al., 2011).

We hypothesized that default network areas would exhibit long-timescale integration properties even in the absence of the hippocampus. We investigated this hypothesis with three primary analyses. 1) *Intact Story Match*: As described above, in neurotypical subjects the activity patterns of default network areas for each paragraph in an intact story depend on the content presented in prior paragraphs. Thus, if the hippocampus is needed for the integration of information across paragraphs, then when amnesic patients listen to an intact story, their default network responses should not match those of controls. Conversely, if default network regions can integrate information across paragraphs without the hippocampus, then amnesic default network activity patterns should match controls when listening to the intact story. 2) *Unscrambling*: Paragraph-scale temporal dependency can be directly tested by comparing amnesic default network responses during the scrambled-paragraphs stimulus to neurotypical default network responses during the intact story, paragraph by paragraph. In neurotypical subjects, default network responses for a given paragraph are changed if the preceding paragraph is changed; if this effect does not depend critically on the hippocampus, then the same should be true in amnesics. In other words, given the long-range temporal dependencies in the default network, the response to the same paragraph should be different between the intact story and the paragraph scrambled conditions for both amnesic and control subjects. 3) *Scramble Reliability:* We additionally examined response reliability of the scrambled-paragraphs stimulus alone, as reduced reliability of scrambled-paragraphs relative to intact story provides complementary evidence of a region’s long-timescale integration properties. As a control test, we predicted that moment-by-moment activity patterns in early auditory areas, which integrate information over short timescales, would be similar between amnesic and neurotypical subjects regardless of paragraph scrambling.

To test these predictions, we used fMRI to record brain responses in neurotypical control subjects and in a bilateral hippocampal amnesic patient as they listened to a 7-minute auditory narrative, as well as to a version of the same stimulus temporally scrambled at the paragraph level. We observed that in some default network regions, including lateral posterior parietal cortex (PPC), posterior medial cortex (PMC), and medial prefrontal cortex (mPFC), there were significantly similar brain activity patterns between the amnesic and controls across the duration of the intact story, suggesting that temporal dependencies at the timescale of paragraphs existed even without the support of the hippocampus. This similarity was observed in spite of the fact that the patient was more than 30 years older than the control subjects—i.e., the 63-year-old amnesic’s brain activity in default network regions substantially resembled that of 18–31-year-old university undergraduate students as they listened to the same real-life story. Furthermore, in both the patient and in the neurotypical control subjects, activity patterns in default network areas for a given paragraph were changed if the preceding paragraph was changed (by scrambling the order of paragraphs), supporting the notion that these brain areas carried information across 30 seconds or more. In contrast, early auditory areas were similar between amnesic and neurotypical subjects regardless of scrambling. This case study provides novel evidence that the purported ability of default network cortical areas to integrate information across long timescales does not depend solely on interactions with the hippocampus.

## Methods

### Participants

An amnesic patient “D. A.” (age: 63 at the time of first fMRI scan session) participated in both behavioral and fMRI tasks. D. A.’s anatomical and behavioral assessments were first reported in Rosenbaum et al. (2008). D. A. became amnesic after contracting herpes encephalitis in 1993. He suffered bilateral MTL damage encompassing the entire right MTL and hippocampus, severe reductions to left MTL cortical areas, and less than ⅓ of the left hippocampus remaining; volume loss was also observed over right-hemisphere posterior temporal, ventral frontal, occipital, and anterior cingulate regions, while left-hemisphere volume loss was limited to the MTL. Small lesions were present in right posterior thalamus and left middle temporal gyrus. See Rosenbaum et al. (2008) for D. A.’s earlier anatomical images, and Figure 1A for new brain images collected for this study in 2015. Behaviorally, he experienced extensive anterograde and graded retrograde amnesia, including memory loss of the period just prior to the onset of amnesia and a postmorbid period, with memories from the most remote time periods spared. His scores on standard neuropsychological tests (Wechsler Memory Scale) are reported in Rosenbaum et al. (2008) and indicate that he has high IQ, preserved short-term memory, and severely impaired delayed memory (<1^st^ percentile). D. A. is a native English speaker with normal hearing and provided written informed consent in accordance with protocols that were approved by the University of Toronto and Baycrest Hospital Research Ethics Boards.

**Figure 1.**
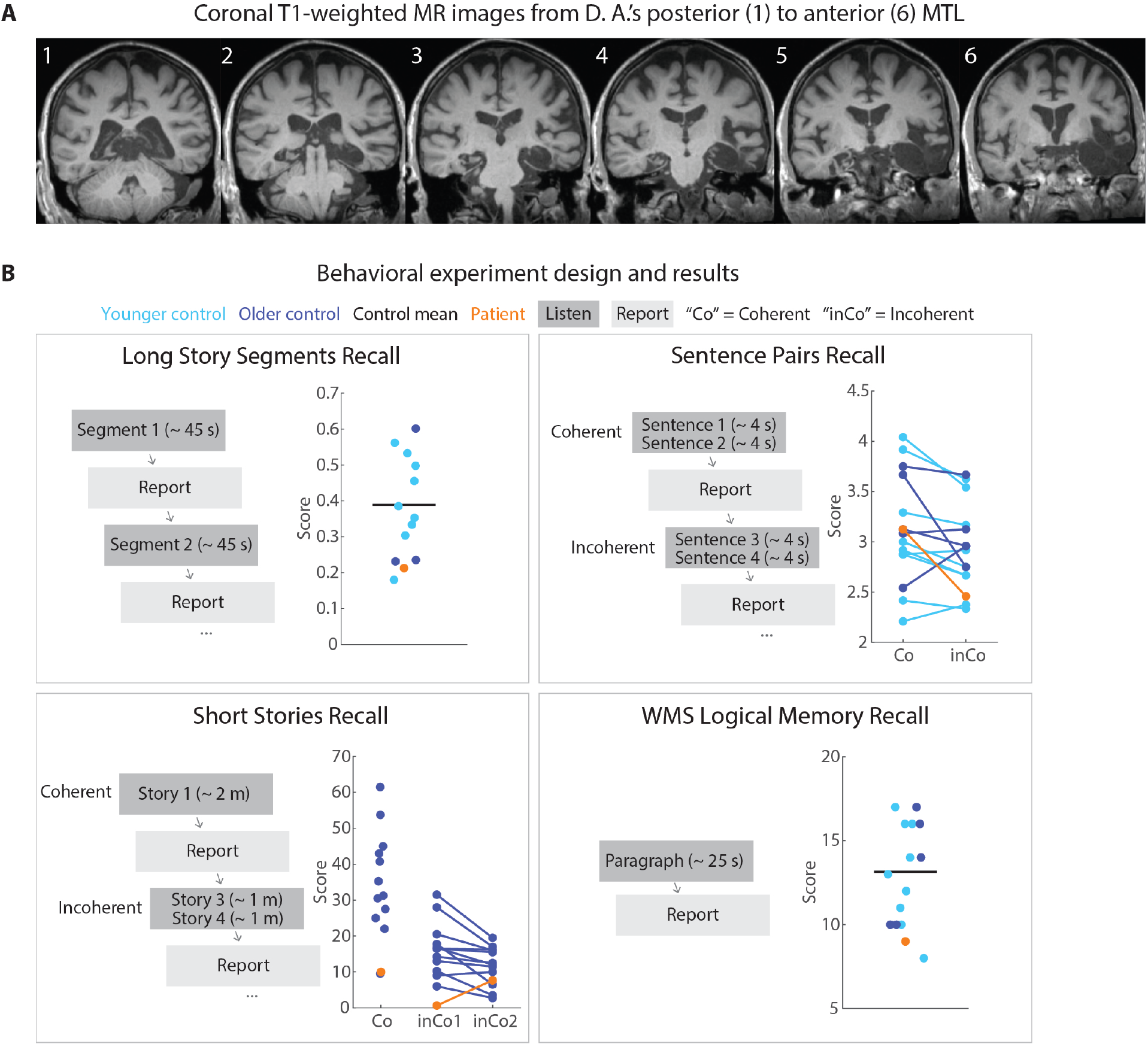
Patient anatomy and behavioral performance on verbal recall tests. **A**) Coronal T1-weighted MR images from D. A.’s posterior to anterior MTL. D. A. has bilateral MTL damage that is more pronounced in the right hemisphere. The right perirhinal, entorhinal, and parahippocampal cortices, and the anterior temporal lobe, are severely damaged. Over 90% of the right hippocampus is damaged. Left perirhinal, entorhinal, and parahippocampal cortices are also severely damaged. **B**) Behavioral task design. Participants listened to four different types of auditory stimuli and verbally reported what they remembered from the stimuli immediately following presentation. In the Long Story Segments session, participants listened to segments (approximately 45-seconds each) from two stories, presented sequentially. Each of the 13 segments (8 from Story 1, 5 from Story 2) was followed by immediate verbal recall. In the Sentence Pairs session, participants listened to 12 coherent (Co) sentence pairs sudo randomly interleaved with 12 incoherent (inCo) sentence pairs, each pair lasting for approximately 4 seconds. Immediately following each sentence pair, participants were instructed to report the sentences verbatim. In the Short Stories session, participants listened to 4 coherent (Co) stories interleaved with 4 incoherent (inCo) short stories, each lasting approximately 2 minutes. The incoherent stories were created by concatenating two different stories. In the WMS-IV Logical Memory experiment, participants listened to the “Anna Thompson” narrative (25 seconds) and reported back the story verbatim.

Age-matched control participants (N = 12, age range: 57–66), as well as a group of younger control participants (N = 9, age range: 18–24; see Table 1 for detailed demographic information), were recruited for behavioral tasks.

**Table 1.** Behavioral experiment control participants

| Session | Older adults | Younger adults | Overall |
| --- | --- | --- | --- |
| Long Story Segments | n = 3 (M age = 63.7; range = 63–64) | n = 9 (M age = 19.2; range = 18–24) | n = 12<br>7 females, 5 males |
| Sentence Pairs | n = 5 (M age = 62.6; range = 67–65) | n = 9 (M age = 19.2; range = 18–24) | n = 14<br>9 females, 5 males |
| Short Stories | n = 12 (M age = 62.8; range = 57–66) | N. A. | n = 12<br>8 females, 4 males |
| WMS | n = 5 (M age = 62.6; range = 67–65) | n = 9 (M age = 19.2; range = 18–24) | n = 14<br>9 females, 5 males |

Control data for fMRI tasks came from a previous experiment, Simony et al. (2016). Thirty-six healthy participants (25 females, ages: 18–33) contributed to the Intact Story condition, and Eighteen participants (12 females, ages: 18–31) contributed to the Scrambled Paragraphs condition. All subjects were native English speakers with normal hearing and provided written informed consent. See Simony et al. (2016) for more details.

### Behavioral tasks: Immediate verbal recall

#### Task design

D. A. participated in four immediate verbal recall tasks: Long Story Segments, Sentence Pairs, Short Stories, and the Wechsler Memory Scale-IV Logical Memory Test (WMS, (Wechsler, 1987). Age-matched controls and younger controls participated in the same four tasks (Figure 1B). For D. A., all audio presentation was controlled by the experimenter and recall was verbally elicited by the experimenter, while control participants used a keyboard to self-initiate listening and recall for each trial.

##### Long Story Segments

Two “long stories” were used, 5 and 8 minutes long, each split into approximately 1 minute segments. The segments were presented sequentially; all participants (including the patient) were instructed to listen to each segment, and then to immediately (within a few seconds) verbally recall that segment only. Participants were instructed to repeat what they had heard, not necessarily verbatim but in as much detail as possible.

##### Sentence Pairs

The sentence stimuli were composed of 12 “coherent” and 12 “incoherent” pairs of sentences. Coherent pairs of sentences shared a continuous context (e.g., both sentences would describe a continuous event happening in a classroom), whereas incoherent pairs of sentences have discontinuous contexts (e.g., one sentence might describe an event taking place on the ocean whereas the other sentence might describe an event happening in the subway). Participants listened to one pair of sentences at a time, and were instructed to repeat what they had heard verbatim. All sentences were between 8 and 13 words; for Coherent pairs, mean lengths were 10.3 words for the first sentence and 10.8 for the second sentence; for Incoherent pairs, mean lengths were 10.6 words for the first sentence (*inCo1*) and 11.2 for the second sentence (*inCo2*). See **Supplementary Materials** for the complete set of sentences.

##### Short Stories

The short story stimuli were composed of 4 “coherent” and 4 “incoherent” short stories. Each coherent story was a 2-minute excerpt from a single story; each incoherent short story was generated by concatenating two 1-minute excerpts from two unrelated stories. For one of the incoherent stories, the first-half story was a repetition of the first-half of one of the coherent stories. All short stories were narrated by the same person (from the podcast *Escape Pod*; see **Supplementary Table 1** for details). Participants were instructed to repeat what they had heard, not necessarily verbatim but in as much detail as possible.

##### Wechsler Memory Scale-IV Logical Memory Test

A narrative paragraph from WMS (*Anna Thompson*, 65 words) was used. Participants listened to the paragraph and were instructed to repeat what they had heard verbatim.

See **Supplementary Table 2** for detailed instructions for all tasks.

#### Scoring

Each participant’s verbal recall was scored using a standardized protocol.

##### Long Story Segments

For each segment, memoranda (“details”) were defined by the experimenters for scoring purposes. The number of details recalled was recorded for each participant. The percentage of number of details recalled out of the total number of details was calculated for each participant.

##### Sentence Pairs

Each recall was given a score from 1 to 5 (5 = produced verbatim; 4 = 1–2 minor semantic and/or syntactic errors; 3 = a few semantic and/or syntactic errors; 2 = a trace of gist; 1 = complete failure).

##### Short Stories

For each short story, memoranda (“details”) were defined by the experimenters for scoring purposes (See **Supplementary Materials**). Transcriptions of participants’ recall were also broken into details. Each recalled detail was given a score (0–2) based on its degree of semantic overlap with the list of details in the original story (2 = complete semantic overlap; 1 = partial semantic overlap; 0 = no semantic overlap, repetitions, metacognitive statements, commentary, and confabulations). The final score was the sum of the scores for all the statements in the recall.

##### WMS

The WMS was scored in the manner prescribed by the scale. There are 25 pieces of information in the paragraph; a point was given for each piece of information recalled accurately.

### fMRI tasks: Auditory narrative listening

Auditory stimuli used for this study were generated from a 7-minute real life story (*Pieman* narrated by Jim O’Grady, recorded at *The Moth*). In the Intact Story condition, participants listened to *Pieman* from beginning to end. In the Scrambled Paragraphs condition, *Pieman* was manually segmented into 12 paragraphs (mean duration 33.9 seconds; s.d. 20.6 seconds) and randomly temporally re-ordered to create the stimulus. Both the Intact Story and Scrambled Paragraphs stimuli were preceded by 12 seconds of music plus 3 seconds of silence, and followed by 15 seconds of silence, all of which were discarded from analyses. For control participants, attentive listening to the story was confirmed using a questionnaire after the scan. See Simony et al (2016) for further details.

D. A. was scanned twice while listening to the *Pieman* Intact Story (data collected in 2015), and twice while listening to the Scrambled Paragraphs stimulus (data collected in 2018). All four scans were used in the analyses.

### MRI acquisition

Control participants, data from Simony et al. (2016), were scanned at Princeton University in a 3 Tesla full-body MRI scanner (Skyra; Siemens) with a 16-channel head coil. Functional images were acquired using a T2* weighted echo planar imaging pulse sequence (repetition time (TR) = 1500 ms; echo time (TE) = 28 ms; flip angle = 64°). Each volume comprised 27 slices of 4 mm thickness (in-plane resolution = 3 × 3 mm^2^; field of view (FOV) = 192 × 192 mm^2^). Slice acquisition order was interleaved. Anatomical images were acquired using a T1-weighted magnetization-prepared rapid-acquisition gradient echo (MP-RAGE) pulse sequence (TR = 2300 ms; TE = 3.08 ms; flip angle = 9°; resolution = 0.89 mm^3^; FOV = 256 × 256 mm^2^). All participants’ heads were stabilized with foam padding to minimize movement. Stimuli were presented using MATLAB Psychophysics Toolbox. Participants were provided with MRI compatible in-ear mono earbuds (Sensimetrics Model S14) to deliver the same audio input to each ear. MRI-safe passive noise-cancelling headphones were placed over the earbuds to attenuate the scanner noise.

The amnesic patient was scanned in two separate sessions. The Intact Story data were collected in 2015 at Rotman Research Institute a 3 Tesla full-body MRI scanner (Siemens). Functional images were acquired using a T2* weighted echo planar imaging pulse sequence (repetition time (TR) = 1500 ms; echo time (TE) = 30 ms; flip angle = 75°). Each volume comprised 28 slices of 4 mm thickness (in-plane resolution = 3 × 3 mm^2^; field of view (FOV) = 192 × 192 mm^2^). Anatomical images were acquired using a T1-weighted magnetization-prepared rapid-acquisition gradient echo (MP-RAGE) pulse sequence (TR = 2000 ms; TE = 2.63 ms; resolution = 1 x 1 mm^3^; FOV = 256 × 256 mm^2^; 160 slices).

The Scrambled Paragraphs data were collected in 2018 at University of Toronto in a 3 Tesla full-body MRI scanner (Prisma; Siemens) with a 20-channel head coil. Functional images were acquired using a T2* weighted echo planar imaging pulse sequence (repetition time (TR) = 1500 ms; echo time (TE) = 30 ms; flip angle = 75°). Each volume comprised 28 slices of 4 mm thickness (in-plane resolution = 3 × 3 mm^2^; field of view (FOV) = 192 × 192 mm^2^).

The patient’s head was stabilized with foam padding to minimize movement. Stimuli were presented using E-Prime. The patient was provided with MRI compatible in-ear mono earbuds (Sensimetrics Model S14) to deliver the same audio input to each ear. MRI-safe passive noise-cancelling headphones were placed over the earbuds to attenuate the scanner noise.

### fMRI preprocessing

Control subjects’ functional data were preprocessed and analyzed using FSL (www.fmrib.ox.ac.uk/fsl), including head motion and slice-acquisition time correction, spatial smoothing (6 mm FWHM Gaussian kernel), and high-pass temporal filtering (140 s period). Preprocessed data were transformed to a standard anatomical (MNI152) brain using FLIRT, and interpolated to 3-mm isotropic voxels.

Patient functional data were preprocessed using the same parameters as controls, with the following modifications. A patient lesion mask was created to identify the lesioned regions for exclusion from calculations of transformation to standard space, and from later analysis (**Supplementary Figure 2**). The mask was manually drawn on D. A.’s MPRAGE anatomical image in native space. It was drawn liberally to exclude all affected areas. The lesion mask was used in conjunction with FSL nonlinear registration tools (FNIRT with option to ignore the masked region) to transform the patient’s anatomical and functional data to standard MNI space and interpolated to 3-mm isotropic voxels.

### Parcellation pattern similarity maps

We computed spatial pattern inter-subject correlation (pISC) (Chen et al., 2017; Nastase et al., 2019) in the Intact Story condition for each of 400 parcels from an independent whole-brain resting-state parcellation (Schaefer et al., 2018). Controls-vs-controls pISC was calculated in the following way. At each time point, the Pearson product-moment correlation coefficient was calculated between 1) one subject’s BOLD activity spatial pattern (i.e., the vector of all voxel values) of a given parcel and 2) the BOLD activity spatial pattern averaged across all other subjects in the same parcel. This process was repeated for all subjects in a given condition and the resulting values were averaged across subjects and across time points to yield a single pISC value (the mean of the diagonal of the time-time correlation matrix). The controls-vs-controls Intact Story pISC values were plotted on the brain surface for every parcel (Figure 2A) using NeuroElf (http://neuroelf.net). The map is displayed at a threshold of r = 0.07 for visualization purposes.

**Figure 2.**
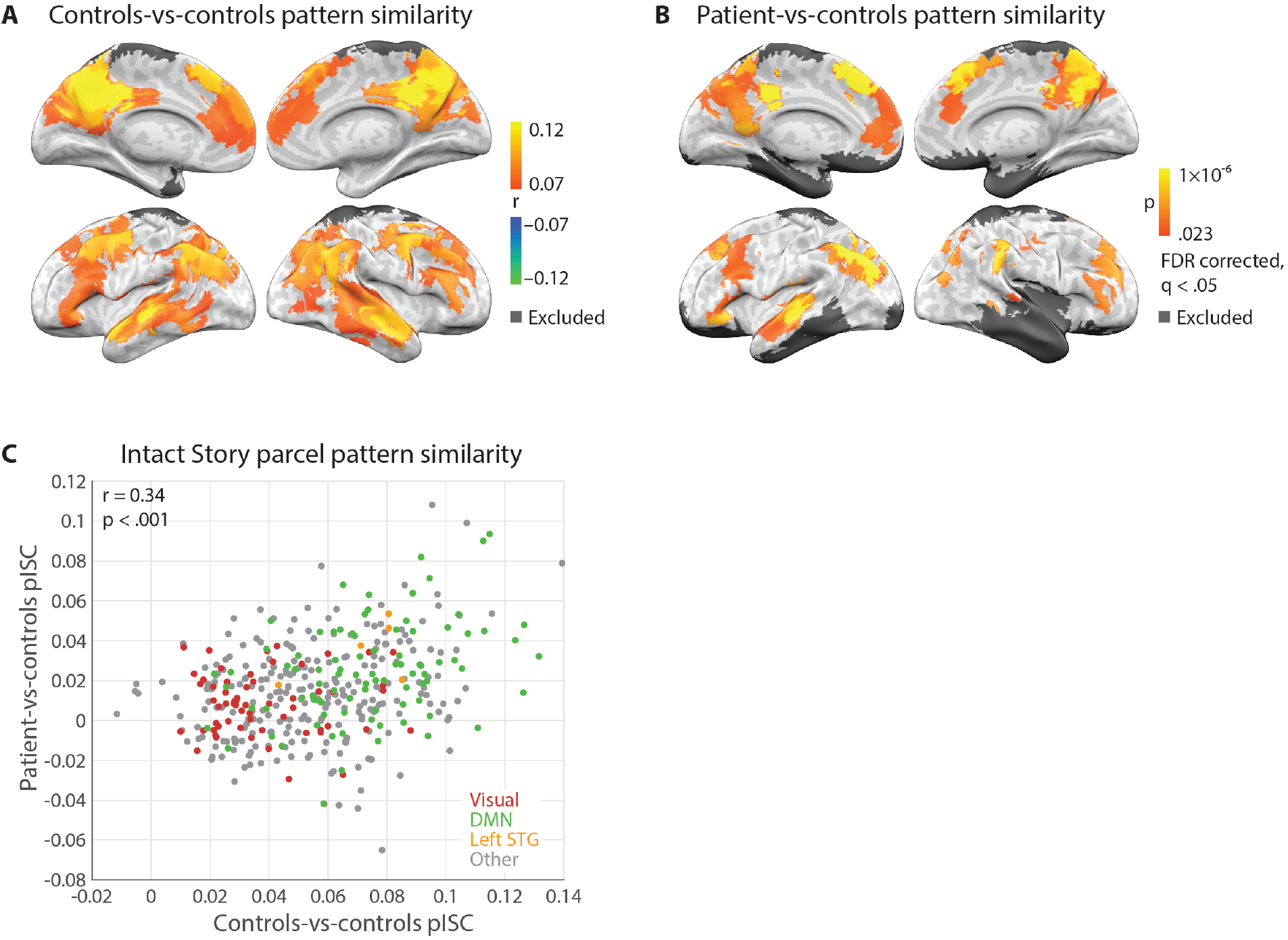
Whole brain spatial pattern inter-subject correlation (pISC) maps of controls-vs-controls and patient-vs-controls in the Intact Story condition. **A**) Whole brain map showing regions where the highest within-group pattern similarity was observed among 36 control subjects, i.e., response reliability map. Threshold r = 0.07 for visualization purposes. **B**) Whole brain map showing regions where significantly similar pattern similarity was observed between the patient and control subjects; one-tailed p-values derived from permutation test and corrected for multiple comparisons across parcels using FDR with q criterion = 0.05. **C**) Across parcels, pattern similarity distribution of patient-vs-controls was similar to that of controls-vs-controls (r = 0.34, p = 5.6 × 10^−12^).

We also calculated patient-vs-controls pISC for Intact Story by correlating, for every timepoint, each of the two patient functional runs with the average of N−1 control subjects, iterating over all possible combinations of N-1 control subjects and then averaging across the N = 36 combinations and across the two functional runs. This procedure matched the controls-vs-controls pISC procedure in that correlations are always calculated between one brain (the patient) and N−1 others (36−1 = 35 controls for the Intact Story condition). Null distributions were generated by randomly shuffling TR-by-TR pattern correlation matrices 10,000 times and retaining the mean diagonal value of each matrix (Kriegeskorte, 2008). As there were many parcels for which no null values exceeded the true pISC value, p-values (one-tailed) were estimated for all parcels by using the null distribution’s mean and standard deviation to fit a normal distribution. We corrected for multiple comparisons across parcels by controlling the False Discovery Rate (FDR) (Benjamini and Hochberg, 1995) using q criterion = 0.05 (Figure 2B). As the test of the patient’s match to controls is predicated on there being an adequate signal in the control data, the patient-vs-controls map is masked with the pISC map of controls-vs-controls (r > 0.07).

We defined a BOLD threshold for excluding parcels with insufficient data. The threshold was set by creating a mean functional image across all conditions and all control subjects and identifying the approximate value (in this case 5000; this threshold may differ for different MRI machines or sequences) above which all remaining voxels fell within the brain. Voxels with values below this threshold, even though they might be within the brain, would be of no greater luminance than voxels falling within the skull or even outside of the head or body, and thus could reasonably be considered regions of BOLD signal dropout. A control subject’s data were retained for a given parcel if at least 50% of voxels in the parcel had mean (of the entire functional run) values above the threshold. As a result, some parcels, especially those near the edge of the brain and signal dropout regions, have a different number of control subjects contributing data. In control-vs-control pISC analyses, a parcel was retained if more than 50% of subjects’ data were retained in that parcel (i.e., more than 18 subjects in the Intact Story condition). Of the 400 original parcels, 296 retained data from 100% of subjects and 387 retained data from more than 50% of subjects; thus 387 parcels were retained for control-vs-control maps. In patient-vs-control pISC analyses, a parcel was retained if 1) more than 50% of control subjects’ data were retained in that parcel; 2) at least 50% of voxels in the average patient data (average of 2 Intact Story functional runs and 2 Scrambled Paragraphs functional runs) pass the retention threshold; and 3) no more than 50% of the voxels in that parcel fell inside the experimenter-defined patient lesion mask. 349 parcels were retained for patient-vs-control maps.

### Regions of interest (ROI) pattern similarity

Regions of interest (ROIs) were created by combining subsets of the parcels: left early auditory cortex, left superior temporal gyrus (STG), left posterior lateral parietal cortex (PPC), left posterior medial cortex (PMC), combined left medial prefrontal cortex (mPFC) and left dorsal medial prefrontal cortex (dmPFC), as well as combined right parahippocampal cortex (PHC) and temporal pole (see **Supplementary Table 3** for a list of parcels used to create these ROIs).

pISC in these ROIs is shown in Figure 3A. Patient-vs-controls pISC and controls-vs-controls pISC were calculated in exactly the same manner as described above for parcels. Null distributions were generated by randomly shuffling TR-by-TR pattern correlation matrices (see **Supplementary Figure 3** 10,000 times and retaining the mean diagonal value of each matrix (Kriegeskorte 2008). The mean diagonal values in the null were plotted as histograms for each ROI. The pISC values was defined as the mean of the diagonal of the original pattern correlation matrix (pISC is plotted as a filled circle).

**Figure 3.**
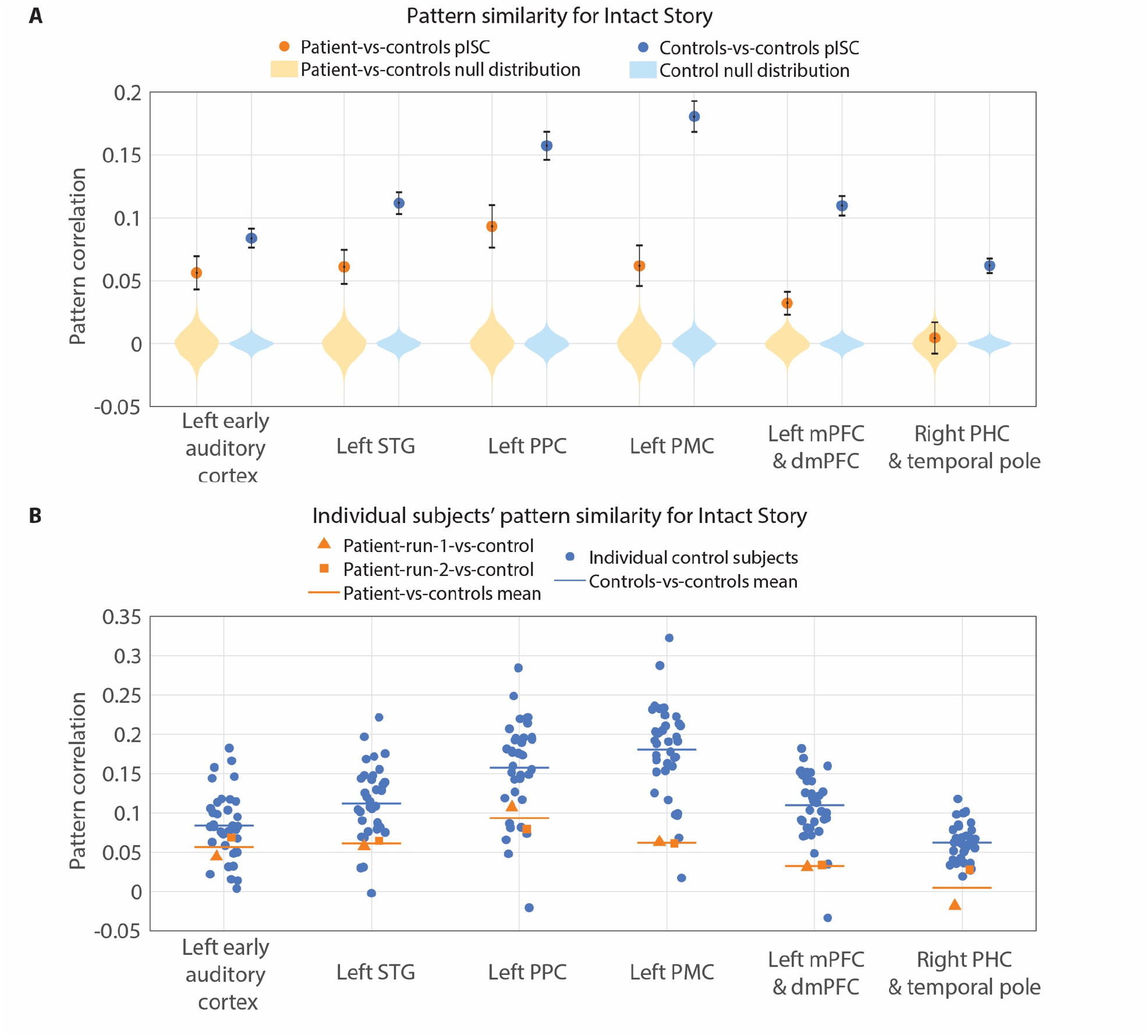
Intact Story pattern similarity in ROIs. **A**) Intact Story spatial pattern inter-subject correlation (pISC) in left auditory cortex, left posterior lateral parietal cortex (PPC), left posterior medial cortex (PMC), combined left medial prefrontal cortex (mPFC) and dorsal medial prefrontal cortex (dmPFC), as well as combined right parahippocampal cortex (PHC) and temporal pole ROIs. Null distributions are generated by randomly shuffling TR-by-TR pattern correlation matrices 10,000 times and retaining the mean diagonal value of each matrix. The mean diagonal values in the null are plotted as histograms for each ROI. The mean diagonal of the original pattern correlation matrix is the true pISC value (plotted as a filled circle). **B**) Individual subjects’ pISC in each ROI.

Measurement error of the pISC values was computed using a bootstrapping approach. We generated a surrogate distribution of the mean pISC values containing 10,000 surrogate diagonals of the time-time pISC matrix. Each surrogate set of pISC values was generated by sampling the pISC values along the diagonal with replacement. The sampling was performed in blocks of 8 contiguous TRs (12 s) so that the surrogate distribution would preserve the temporal autocorrelation induced by the hemodynamics. After each surrogate set of pISC values was computed, the mean of that surrogate set was computed across all timepoints. The standard deviation of this distribution then provided the standard error of the mean (error bar on filled circles in Figure 3A).

Comparison of pISC between groups was performed with two-tailed unpaired t-tests of the 36 patient-vs-controls to the 36 controls-vs-controls values, separately for early auditory cortex and STG, for Intact Story. Comparison of patient-vs-controls pISC between mPFC and PPC, and between mPFC and PMC, was performed with two-tailed unpaired t-tests of the 36 patient-vs-controls values for each ROI.

Individual control subject and individual patient functional run pISC values are shown in Figure 3B.

### Unscrambling analyses

In order to compare Intact Story brain responses to Scrambled Paragraphs brain responses, it was necessary to re-order the Scrambled Paragraphs data such that the acoustic input would be matched at each timepoint across the two conditions. In other words, the Scrambled Paragraphs needed to be “unscrambled” to reconstruct the same ordering from the Intact Story.

We first manually identified (using Adobe Audition) the onset and offset of speech in the Intact Story and Scrambled Paragraphs stimuli, thus excluding pre-story music and post-story silence. We then identified boundaries of the 11 paragraphs in the Scrambled Paragraphs stimulus and the same boundaries in the original Intact Story stimulus. All boundaries were recorded with a temporal resolution of 44.1 KHz (0.002 seconds). As the BOLD data were collected at a 1.5-second TR resolution, paragraph boundaries did not necessarily fall at the beginning of a TR. To avoid the accumulation of small errors during the unscrambling procedure, we resampled the fMRI time courses to a higher temporal resolution (50 Hz) to match the Intact and the Scrambled Paragraphs fMRI time courses more precisely. To verify the beginning and end times of the auditory stimulus in the fMRI time course, the correlation was calculated between 1) each participant’s fMRI time course in the auditory cortex and 2) the audio envelope of the stimulus (both resampled to 50 Hz) at all possible lags to find the peak correlation. This procedure also estimates the magnitude of delay due to the hemodynamic response function: in this case the cross-correlated peaked at a lag of 3 TRs = 4.5 seconds. Thus, the fMRI time course was cropped and shifted by 3 TRs. We then “unscrambled” the fMRI time course of the Scrambled Paragraphs condition to match the temporal order of the Intact Story condition. The data (Intact Story and now “Unscrambled Paragraphs” conditions) were resampled back to the original frequency (1/1.5 Hz) for all subsequent analyses.

pISC was calculated between Intact Story and Unscrambled Paragraphs conditions for controls-vs-controls and patient-vs-controls for every parcel falling within the controls-vs-controls Intact Story reliability map (r > 0.07, Figure 2A). Null distributions were generated by randomly shuffling TR-by-TR pattern correlation matrices 10,000 times and retaining the mean diagonal value of each matrix (Kriegeskorte, 2008). One-tailed p-values are plotted for each brain parcel in Figure 4A–B. The maps were masked with controls-vs-controls Intact Story reliability (pISC) r > 0.07, and FDR correction was performed within this mask.

**Figure 4.**
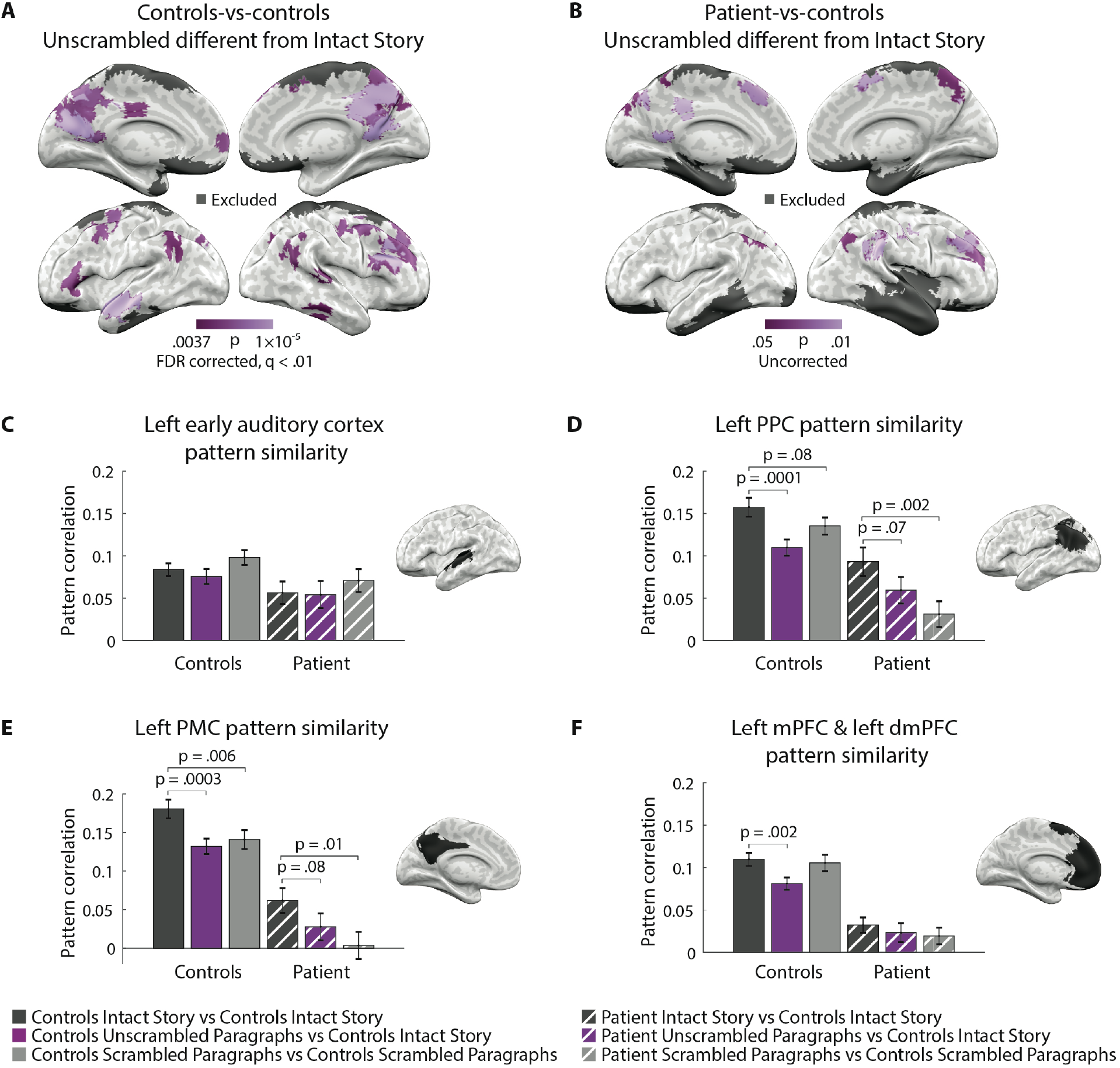
Unscrambling analysis. **A**) Parcels for which pattern correlation of controls-vs-controls Intact Story was higher (p < 0.01) than controls Intact Story vs. controls Unscrambled Paragraphs. Map masked with controls-vs-controls Intact Story reliability (pISC) r > 0.07. **B**) Parcels for which pattern correlation of patient-vs-controls Intact Story was higher (p < 0.05) than patient Unscrambled Paragraphs vs. controls Intact Story. Map masked with controls-vs-controls Intact Story reliability (pISC) r > 0.07. **C–F**) Pattern similarity for Intact Story (dark grey), Intact Story vs. Unscrambled Paragraphs (purple), and Scrambled Paragraphs (light grey); controls-vs-controls (solid) and patient-vs-controls (striped); in left posterior lateral parietal cortex PPC, left posterior medial cortex (PMC), left medial prefrontal cortex (mPFC), and left early auditory cortex.

pISC was calculated between Intact Story and Unscrambled Paragraphs conditions for controls-vs-controls and patient-vs-controls in *a priori* ROIs: left PPC, left PMC, left mPFC, and left early auditory cortex (Figure 4C–F). pISC measurement error was calculated using the same block bootstrap approach as for Figure 3A. Between-condition comparisons of pISC in Figure 4C–F were calculated by generating surrogate distributions of the difference between conditions, using a bootstrapping approach (sampling 10,000 times with replacement) and the same blocking parameters as above. One-tailed p-values are reported.

### Patterns across time

Controls-vs-controls and patient-vs-controls pISC were examined at individual timepoints and time bins across the duration of the Intact Story. TR-by-TR pISC is displayed for the left auditory cortex and left PPC ROIs (Figure 5A & 5B). For coarser time bin analysis, bins were created by evenly dividing the Intact Story into three segments and averaging pISC across time within each segment (Figure 5C & 5D, left panels). pISC measurement error for the segments was calculated using the same block bootstrap approach as for Figure 3A. Comparison of whether the difference between controls-vs-controls and patient-vs-controls pISC changed across the segments was conducted with two-tailed unpaired t-tests of the difference.

**Figure 5.**
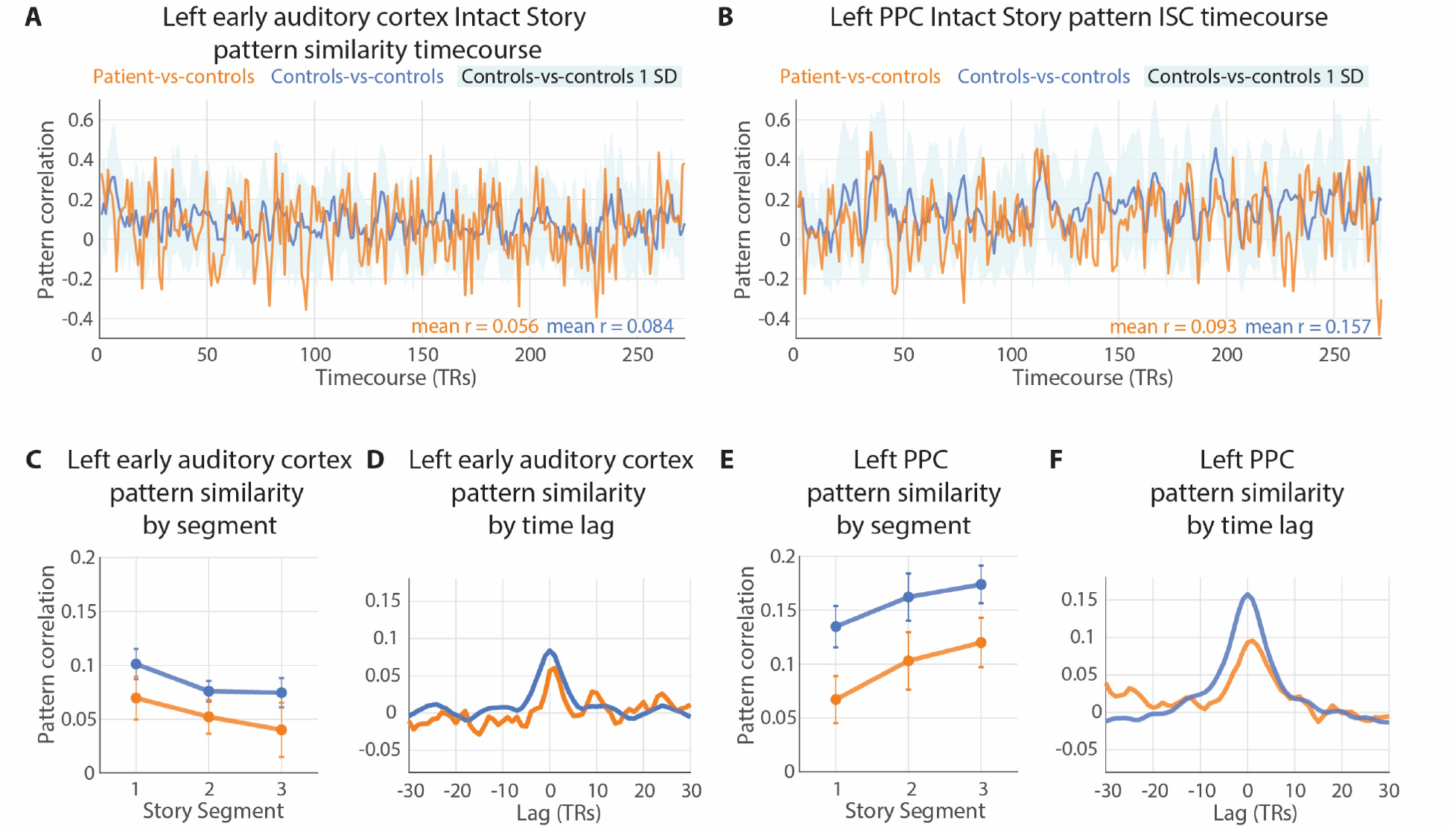
Pattern similarity across time. **A** & **B**) Patient-vs-controls and controls-vs-controls spatial pattern inter-subject correlation in left auditory cortex and left posterior lateral parietal cortex (PPC) at every timepoint of the Intact Story. Standard deviation of controls-vs-controls at each timepoint plotted in light blue. Mean values of each pattern similarity timecourse printed in lower right corner. **C & E**) The Intact Story data were evenly divided into three segments and averaged across time within each segment. Patient-vs-control and control within-group pISC were computed for each segment. **D & F**) pattern similarity lag correlation of patient-vs-controls and controls-vs-controls in auditory cortex and PPC. The peaks at lag zero, with gradual declines as lag magnitude increases (either negative or positive), indicate that the pattern match between patient-vs-controls (orange) and controls-vs-controls (blue) was temporally specific, i.e., dependent on the stimulus matching moment-by-moment.

To examine the temporal specificity of pISC, we computed voxel pattern lag correlation timecourses by calculating the pISC of controls-vs-controls and of patient-vs-controls at every possible temporal offset (e.g., time lag = 1 TR, 2 TRs, etc.) from −30 to +30 TRs (Figure 5C & 5D, right panels). This procedure requires that TRs with no corresponding data between conditions be removed from analysis; e.g., at lag = +30 TRs, the patient data and control data are offset by 30 TRs, and thus 30 TRs are dropped from the beginning of one data series and another 30 TRs are dropped from the end of the other data series.

### Pattern vs. temporal ISC

Spatial pattern inter-subject correlation (pISC) and temporal inter-subject correlation (tISC) were calculated for every parcel in the brain using the Intact Story data, both for controls-vs-controls and for patient-vs-controls, and visualized in scatter plots (**Supplementary Figure 4**). Visual and default mode networks refer to a standard 7-Network resting-state parcellation (Schaefer et al., 2018; Yeo et al., 2011). Auditory parcel selection is described above in *Methods: Regions of interest*; see also **Supplementary Table 3**.

## Results

### D. A. displays behavioral patterns of anterograde amnesia in tests of verbal recall

As the goals of the current experiment are to investigate default network activity in the absence of a functioning hippocampus, we first sought to establish that D.A. displays behavioral patterns concomitant with severe bilateral hippocampal damage, i.e., anterograde amnesia: impaired but not absent immediate prose recall, catastrophic memory loss after a filled delay, performance disruption from discontinuity (e.g., topic changes), and preserved short-term memory when active rehearsal is allowed (Squire and Wixted, 2011). Rosenbaum et al. (2008) previously reported that D. A. has dense anterograde amnesia, e.g., <1^st^ percentile on delayed memory tasks in the Wechsler Memory Scale (WMS) (Wechsler, 1987). Here we extend these prior findings with additional tests of immediate verbal recall. To probe his comprehension of semantically-rich material, we constructed tests using sentences and narratives of different lengths and coherence. D. A. and controls participated in four immediate verbal recall tasks: WMS, Long Story Segments, Short Stories, and Sentence Pairs (Figure 1B).

The WMS Logical Memory test (Wechsler, 1987) is a common measure of “prose” recall in amnesic patients (e.g., Baddeley and Wilson, 2002; Kopelman, 1987; Rosenbaum et al., 2008). In this test the experimenter reads aloud a 65-word story, and the patient is then asked to repeat it verbatim. D. A.’s recalled 9 out of a possible 24 items from the story Figure 1B). This value was lower than all of the age-matched controls, second-lowest amongst the younger controls, and is typical for amnesic individuals, (range of 0–15 memory units for the population tested by Baddeley and Wilson, 2002).

To test the limits of more natural and extended narrative comprehension in amnesia, we also measured D.A.’s verbal recall for longer narratives. In the Long Story Segments task, auditory stories of 5 and 8 minutes were played 1 minute at a time. After each such segment, all participants were asked to repeat what they had heard, not necessarily verbatim but in as much detail as possible. D. A.’s performance on this test was poor but within the range of controls (second-lowest; Figure 1B).

We next examined narrative recall while manipulating the coherence or incoherence of the narrative. In the Short Stories task, “coherent” auditory stories were created by excerpting 2 minutes from a single story (*Co*), and “incoherent” stories generated by concatenating two 1-minute excerpts from two unrelated stories (*inCo1* and *inCo2*). Again, the patient and controls were asked to repeat what they had heard, not necessarily verbatim but in as much detail as possible. D. A.’s performance on this test was poor (lowest in both conditions; Figure 1B). Notably, when incoherent stories were scored separately for the first and second half (*inCo1* and *inCo2*), D. A.’s performance was near the mean of controls for the second halves of stories, but he recalled only one gist-like detail from the first half of any of the incoherent stories. Here, the discontinuity introduced by the concatenation of two different stories may have impacted D.A.’s ability to retrieve the material from before the discontinuity (event boundary) that occurred one minute earlier. Unfortunately, it was not possible to score coherent stories separately for the first and second half, as D.A.’s (and many controls subjects’) verbal reports were largely gist-like, and thus very few of the recall statements could be clearly identified as coming from the first or second half of the coherent story.

We also assessed recall for shorter linguistic sequences. On each trial of the Sentence Pairs task, participants listened to two consecutive sentences (each sentence composed of 8-13 words) and then immediately attempted to repeat verbatim what they had heard. D.A.’s performance was close to the mean of controls for “coherent” sentence pairs (when the two sentences described a continuous context) and poor (3^rd^ lowest) for “incoherent” sentence pairs (when the two sentences described different contexts; Figure 1B). These results converge with studies of hippocampal amnesics showing that prior knowledge can improve short-term memory retention for sentences (Race et al., 2015b).

After the first scan session (described below), and several minutes after D. A. had exited the MRI machine, the experimenter asked D. A. whether he remembered anything from the auditory narrative he had listened to inside the scanner. D. A. could not verbalize any accurate memory for content from the story; he made only statements that were either so vague as to be indistinguishable from guesses, or completely incorrect, even when prompted with several story details (see **Supplementary Materials** for a transcript of the interview).

Taken together, D. A.’s behavior patterns match those of anterograde amnesic syndrome arising from severe bilateral hippocampal damage. He demonstrated preserved immediate recall of short sentences (coherent Sentence Pairs), and impaired recall of even very recent (a few seconds ago) material prior to discontinuities introduced by an unrelated sentence (incoherent Sentence Pairs). His immediate verbal recall performance was poor but within the range of control participants. For two tests, 1) recall of coherent Sentence Pairs, and 2) recalling the second half of an Incoherent Story, his performance was close to the mean of controls. D.A.’s prose and narrative immediate recall were poor but not completely absent (WMS, Long Story Segments, coherent Short Stories); and he had virtually no memory of events prior to a discontinuity one minute earlier (incoherent Short Stories).

### D. A.’s neural responses to an auditory narrative match the responses of controls (Intact Story Match)

The primary question of the study was whether cortical regions of the default network would exhibit long-timescale properties even in the absence of a hippocampus. In neurotypical subjects the activity patterns of default network areas for each paragraph in a continuous story depend on the content presented in prior paragraphs (Hasson et al., 2015; Lerner et al., 2011). If information from the prior paragraph is not being carried forward into the current paragraph in the patient brain, the patient’s default network activity pattern for each paragraph should be different from the neurotypical pattern. In other words, if default network regions can integrate information across paragraphs without the hippocampus, then amnesic default network activity patterns should match controls when listening to the intact story. To test this, we compared brain activity between the patient and control participants as they listened to the same intact 7-minute auditory narrative. (Note that the patient does have residual hippocampal tissue— less than ⅓ of the left hippocampus remains—but 1) he presents behaviorally as densely amnesic, and 2) meta-analysis of non-human primate data shows that partial hippocampal loss can be equivalent or worse to more severe damage in terms of its impact on delayed memory performance (Baxter and Murray, 2001).)

First, we established the regions where control participants exhibited similar brain activity patterns with each other while listening to the intact narrative by calculating pattern similarity in every parcel (Schaefer et al., 2018) across the brain. Previous fMRI studies of the temporal integration hierarchy have largely used temporal ISC (tISC; e.g., Hasson et al., 2008; Lerner et al., 2011; Simony et al., 2016); here we use spatial pattern ISC (pISC; e.g., Chen et al., 2017; Oedekoven et al., 2017; Baldassano et al., 2018; Nastase et al., 2019) which allows estimates of response reliability to be calculated at each TR. The two are closely related but not redundant ((Nastase et al., 2019); **Supplementary Figure 4**). We calculated pattern similarity for controls-vs-controls at every TR of the Intact Story and averaged across all TRs within each parcel (Figure 2A). In agreement with prior studies, the spatial patterns of neural responses were similar between control subjects in widespread regions, ranging from early auditory areas, to linguistic areas, to default network areas including posterior medial cortex (PMC), lateral posterior parietal cortex (PPC), and medial prefrontal cortex (mPFC) (Chen et al., 2017; Baldassano et al., 2018; Nastase et al., 2019). Controls-vs-controls whole-cortex response reliability map (Figure 2A) is shown at r > 0.07 for visualization purposes.

Next, we compared the patient’s moment-by-moment brain activity to that of control participants as they listened to the same Intact Story. Mirroring the analysis above, we calculated pattern similarity for patient-vs-controls at every TR of the Intact Story and averaged across all TRs within each parcel (Figure 2B). Brain responses of the amnesic patient were similar to responses in neurotypical control subjects in many cortical areas, including early auditory areas, bilateral PMC, bilateral PPC, and bilateral mPFC. Note that the patient’s more severe damage is in the right hemisphere temporal lobe. Whole-cortex maps show one-tailed p-values derived from permutation test and corrected for multiple comparisons using FDR with q criterion = 0.05.

To compare controls-vs-controls and patient-vs-controls distributions of pattern similarity across the brain, we visualized pattern similarity of individual parcels in a scatter plot (Figure 2C). Across parcels, the pattern similarity of patient-vs-controls was significantly similar to that of controls-vs-controls (r = 0.34, p < 0.001). We also evaluated this relationship when restricting to a higher threshold, as parcels with very low correlations might be considered noise. Using a threshold of r > 0.07 for controls-vs-controls and r > 0.03 for patient-vs-controls (same thresholds as Figures 2A–B), 55 parcels were retained, and the correlation of pattern similarity across parcels between patient-vs-controls and controls-vs-controls was r = 0.31, p = 0.02).

### Significant similarity of neural patterns between patient and controls in default network ROIs (Intact Story Match)

Our main interest was in default network regions, which were previously shown to integrate information over long timescales during continuous natural input, such as movies and stories (Hasson et al., 2008; Lerner et al., 2011; Simony et al., 2016; Chen et al., 2016). Thus, we examined pattern similarity in three default network ROIs (PPC, PMC, mPFC), as well as in (i) early auditory cortex and (ii) a larger superior temporal gyrus (STG) ROI that included early auditory cortex. As the patient’s brain damage is most severe in the right hemisphere (including volume loss in right-hemisphere cortical areas outside of the MTL), ROI analyses were restricted to the left hemisphere, with the exception of a control region in the right anterior temporal and parahippocampal cortex where there is little tissue remaining. For each ROI, pattern similarity was calculated for controls-vs-controls and for patient-vs-controls at every TR of the Intact Story and averaged across all TRs (same analysis as conducted for each parcel in Figure 2). A null distribution was created for each ROI by randomly permuting TR-by-TR pattern correlation matrices (**Supplementary Figure 3**) and recalculating the mean diagonal 10,000 times.

We expected auditory cortex activity patterns to be similar across patients and controls, as these areas respond primarily to immediate acoustic features of the auditory narrative. This prediction was confirmed: the true patient-vs-controls pattern similarity value was positive and well outside the null distribution for both the early auditory cortex (M = 0.056, p < 0.001) and STG (M = 0.061, p < 0.001); Figure 3A). Controls-vs-controls pattern similarity was also significant in these two regions (early auditory: M = 0.084, p < 0.001; STG: M = 0.11, p < 0.001) and higher than patient-vs-controls pattern similarity (early auditory: t(70) = 3.81, p < 0.001; STG: t(70)=6.49, p < 0.001; Figure 3A). These data suggest that fMRI signal in the control data is more reliable overall, possibly due to the lower age of the control participants (Campbell et al., 2015) or other idiosyncrasies of the patient testing. Nonetheless, the statistically reliable match between patients and controls is positive evidence of a common response between amnesic and neurotypical responses.

In PPC, PMC, and mPFC, default network regions which were previously shown to integrate information over long timescales, we observed similar activity between patient and controls (Figure 3A), with the true correlation values positive and falling well outside of the null distributions (PPC: M = 0.093, p < 0.001; PMC: M = 0.062, p < 0.001; mPFC: M = 0.032, p < 0.001). In mPFC the match between between patient and controls was weaker than in PPC and PMC (mPFC vs. PPC: t(70) = 193.8, p < 0.001; mPFC vs. PMC: t(70) = 103.6, p < 0.001). In all three of these default network ROIs, controls-vs-controls pattern similarity was significantly positive (PPC: M = 0.16, p < 0.001; PMC: M = 0.18, p < 0.001; mPFC: M = 0.11, p < 0.001), as observed in prior studies (Chen et al., 2017; Baldassano et al., 2018).

In right PHC and temporal pole, a severely damaged region of the patient’s brain, the true patient-vs-controls correlation fell inside the null distribution (M = 0.005) i.e., there was no similarity effect. This is as expected given that virtually no BOLD signal was detectable in this area. This ROI normally shows positive pattern similarity between individuals during narrative listening, as demonstrated in the controls-vs-controls comparison (M = 0.062, p < 0.001; Figure 3A).

Data for individual subjects in all ROIs are shown in Figure 3B.

Altogether, the results in Figures 2–3 show that patterns of activity during Intact Story listening were similar between the patient and controls across the brain. This was true for a number of individual parcels and *a priori* default network ROIs, and the overall distribution of patient-vs-controls pattern similarity strength across parcels mirrored that of the control subjects. Because the activity patterns of default network areas for each paragraph in a continuous story depend on the content presented in prior paragraphs in neurotypical subjects (Lerner et al., 2011; Hasson et al., 2015), these results suggest that default network regions can, to some extent, integrate information across paragraphs without the hippocampus. However, the match between patient and controls was weaker overall than the match of controls to each other. While this could be due to the patient being older than controls or other idiosyncrasies of the patient brain, it is also possible that default network patterns reflect a mixture of long-and shorter-timescale information, and the preserved component of the patient’s default network activity pattern corresponded to shorter-timescale information; the Intact Story condition alone cannot conclusively exclude this interpretation. Thus, we examined the Scrambled Paragraphs condition of the experiment.

### Default network responses affected by paragraph scrambling, even in absence of the hippocampus (Unscrambling and Scramble Reliability)

In prior studies, default network regions were shown to be able to integrate information across paragraphs, on the scale of 30 seconds or more, during continuous narrative processing. This was demonstrated in two complementary ways: 1) comparing Intact Story timecourses to “unscrambled” Scrambled Paragraphs timecourses, which showed that if the preceding paragraph was altered, then default network responses to the current paragraph were altered (Lerner et al., 2011); and 2) calculating temporal ISC for a Scrambled Paragraphs narrative, which showed that default network inter-subject response reliability was reduced when paragraph-level (or an equivalent window of time for visual stimuli) information was disrupted (Chen et al., 2016; Hasson et al., 2008; Honey et al., 2012; Lerner et al., 2011; Simony et al., 2016). By contrast, early auditory areas exhibit high inter-subject response reliability regardless of scrambling. We sought to replicate these effects in the current dataset, both in controls and in the patient, by comparing brain responses during Intact Story to those recorded during Scrambled Paragraphs.

First, we “unscrambled” the Scrambled Paragraphs brain data by identifying the TRs corresponding to each paragraph and re-ordering these to match the Intact Story. This created an “Unscrambled Paragraphs” version of the data in which each TR was time-aligned to the Intact Story, i.e., the same moment of the stimulus was presented for each TR of both conditions. Then, for each parcel of the brain, we calculated spatial pattern correlations between Unscrambled Paragraphs and Intact Story and tested whether this value was significantly different from pattern similarity between subjects listening to the Intact Story (the theoretical ceiling of response reliability). The unscrambling analysis confirmed that, in controls, default network regions such as PPC, PMC, and mPFC produced responses that did depend on the content of preceding paragraphs (map FDR corrected at q < 0.01), while low-level regions, e.g., early auditory cortex, produced responses that did not depend on prior paragraphs (Figure 4A). A subset of the same regions was identified in the patient-vs-controls comparison using a less conservative threshold; two parcels passed FDR correction of q < 0.05: one in left PMC, and one in right PPC (157 and 304 in the Schaefer et al. (2018) atlas). The map in Figure 4B is shown uncorrected at p < 0.05 for visualization purposes. In *a priori* ROIs, we observed significant effects of scrambling (unscrambled-vs-intact less correlated than intact-vs-intact) in PPC, PMC, and mPFC for controls-vs-controls (PPC: p = 0.0001; PMC: p = 0.0003; mPFC: p = 0.002); and trends for the same effect in PPC (p = 0.07) and PMC (p = 0.08), but not mPFC (p > 0.1), for patient-vs-controls (Figure 4D–F, dark grey vs. purple bars).

Scrambling did not alter response patterns in early auditory cortex for either controls-vs-controls or for patient-vs-controls (Figure 4C, ps > 0.1; dark grey vs. purple bars). Second, we examined reliability (spatial pattern similarity) in the Scrambled Paragraphs condition for controls-vs-controls and patient-vs-controls. Echoing previous studies using temporal ISC with neurotypical subjects (Lerner et al., 2011; Simony et al., 2016), inter-subject pattern similarity strength in PPC and PMC was reduced by paragraph scrambling in the controls-vs-controls comparisons (PPC: p = 0.08; PMC: p = 0.006); we observed a similar pattern in the amnesic-vs-controls comparisons (PPC, p = 0.002; PMC, p = 0.01); but not in mPFC (Figure 4C–E, dark grey vs. light grey bars). As predicted, inter-subject pattern similarity strength in these auditory areas was unaffected by paragraph scrambling (Figure 4F, dark grey vs. light grey bars). For STG and PHC see **Supplementary Figure 5**. Right hemisphere ROIs are displayed in **Supplementary Figure 5**.

In sum, we observed that scrambling at the level of paragraphs affected the amnesic’s default network responses in two ways: 1) the unscrambling analysis showed that if the preceding paragraph was altered, then default network responses to the current paragraph were altered; and 2) response reliability, measured using pattern similarity at each TR, was reduced when paragraph-level information was disrupted by scrambling. These results support the interpretation that the amnesic’s default network regions were able to integrate information across paragraphs during the intact story, even in the absence of the hippocampus.

### Patient-vs-controls neural pattern similarity is sustained across the duration of the story

While the above analyses examined whether the brain activity of the patient and controls was similar on average across all moments of the Intact Story, it was also important to test whether the similarity persisted across the full duration of the stimulus. Potentially, the patient could resemble controls most strongly at the beginning of the story, with similarity decreasing over time as the patient’s memory of more distant prior information weakened.

Thus, we visualized pattern similarity, TR-by-TR, in left auditory cortex and left PPC (Figure 5A–B). In both regions, more timepoints were positive than negative (as should be expected given the positive means reported in Figure 2A), for both patient-vs-controls (orange bars) and controls-vs-controls (blue lines). Note that, while the correlation magnitudes are comparable between the patient-vs-controls (orange bars) and controls-vs-controls (blue lines), the measurement precision is different across the two curves. This is because patient-vs-controls values reflect an average of 2 samples while the controls-vs-controls values reflect an average of 36 samples. See **Supplementary Figure 6** for timecourses in all *a priori* ROIs.

The similarity between patient and controls did not decrease over time in PPC (Figure 5). We computed the average pattern similarity across the first third, middle third, and final third of the story. In auditory cortex, the correlation values numerically decreased slightly over time in both patient-vs-controls (orange) and controls-vs-controls (blue) (Figure 5C). In any region, such a decrease might be due to either fatigue, or to properties of the stimulus that varied across segments of time—e.g., in auditory cortex, one segment might have lower volume variability than another and thus lower signal to drive ISC. Note that, despite the drop over time in auditory cortex, the *difference* between controls-vs-controls and patient-vs-controls remained constant (ps > 0.3). In PPC, where one might have expected a decrease in patient-vs-controls over the duration of the story, there was no such drop (Figure 5E). Numerically, both patient-vs-controls and controls-vs-controls pattern similarity actually increased across thirds of the story. Again, the *difference* between controls-vs-controls and patient-vs-controls remained constant (ps > 0.4). These results are consistent with the interpretation that the patient’s comprehension of the narrative was similar to controls even toward the later periods of the story.

To confirm that the pISC values reflected time-locked responses to the auditory narrative, we also evaluated the temporal precision of the match between patient and controls. Figure 5D,F show the pattern similarity lag correlation of patient-vs-controls and controls-vs-controls in auditory cortex and PPC. The peaks at lag zero, with gradual declines as lag magnitude increases (either negative or positive), indicate that the pattern match between patient-vs-controls (orange) and controls-vs-controls (blue) was temporally specific, i.e., dependent on the stimulus matching moment-by-moment.

## Discussion

Default network areas are proposed to carry slowly-changing information during continuous semantically-rich input such as stories and conversation (Hasson et al., 2015). In healthy people, these areas are functionally coupled to the hippocampus and may work together to accumulate, maintain, and integrate information across time during naturalistic input. However, it is an open question whether the long-timescale capability of default network regions depends critically on interactions with the hippocampus. In this study we investigated whether default network areas can integrate information over tens of seconds even *without* the hippocampus by recording the brain activity of an amnesic patient with severe bilateral hippocampal damage as he listened to a seven-minute auditory story. We observed that in some default network regions, including lateral posterior parietal cortex (PPC), posterior medial cortex (PMC), and medial prefrontal cortex (mPFC), there were significantly similar brain activity patterns between the amnesic and controls across the duration of the intact story, suggesting that temporal dependencies at the timescale of paragraphs existed even without the support of the hippocampus. Furthermore, in both the patient and in neurotypical subjects, activity patterns in some default network areas (PPC and PMC, but not MPFC) for a given paragraph were changed if the preceding paragraph was changed (by scrambling the order of paragraphs), supporting the notion that these brain areas carried information across 30 seconds or more. In contrast, early auditory areas were similar between amnesic and neurotypical subjects regardless of scrambling. This study provides novel evidence that the purported ability of default network cortical areas to integrate information across long timescales does not depend solely on interactions with the hippocampus.

Prior studies left open the question of whether the long-timescale capability of default network regions depends on interactions with the hippocampus. Initial experiments used parametrically scrambled narratives, both auditory and visual, to show that temporal activity profiles in default network areas were contingent on narrative segments being intact on the scale of 30 seconds or more (Hasson et al., 2008; Lerner et al., 2011; Honey et al., 2012). In Chen et al. (2016), healthy subjects viewed movie events whose comprehension depended on prior information presented either 1) several minutes earlier within the same viewing session (i.e., without major discontinuity) or 2) a day earlier (introducing major discontinuity). The day-earlier condition differentially recruited hippocampus and evinced strong stimulus-locked functional connectivity between the hippocampus and cortical default network areas, with default network responses between the groups initially dissimilar and converging after 3–4 minutes. One interpretation of these results is that hippocampus-default network interaction was important for bridging the discontinuity, and, conversely, that in the continuous condition the default network regions were able to support story comprehension with less or no hippocampal interaction. However, all of these experiments were conducted in neurologically healthy individuals, and thus could not answer the question of whether the hippocampus is necessary for the long-timescale properties of default network regions.

While our results suggest that default network cortical areas are capable of supporting online processing and comprehension of real-world events without the hippocampus, this does not contradict the view that default network-hippocampus interactions are needed for encoding and later retrieval of the event information. Even if their online processing of stimuli is preserved, hippocampal amnesics are nonetheless unable to retrieve episodic information at a later time, either due to disrupted encoding or retrieval. Studies of functional connectivity between the hippocampus/MTL and default network during naturalistic input have emphasized the interaction between these structures in service of memory encoding and retrieval in the healthy brain (van Kesteren et al., 2010; Chen et al., 2016; Bonasia et al., 2018; Aly et al., 2018), and hippocampal responses at the boundaries between events are theorized to support consolidation of event information (Ben-Yakov et al., 2013; Baldassano et al., 2017; Ben-Yakov and Henson, 2018). Interestingly, MTL damage does not impact connectivity among default network nodes during resting state, but does alter connectivity between the MTL and default network areas (Hayes et al., 2012); in fact, it has been proposed that abnormal hippocampal interactions with other brain regions are the cause of behavioral deficits in amnesia, as opposed to hippocampal damage per se (Argyropoulos et al., 2019). In an experiment closely related to the current study, Oedekoven et al. (2019) scanned a patient with anterograde amnesia resulting from thalamic stroke as they viewed video clips and during later cued retrieval. While no univariate activity differences were observed between the patient and controls during video viewing and retrieval, functional connectivity between bilateral PMC and left hippocampus was reduced in the patient brain relative to controls during both video viewing and retrieval. Such results are consistent with the notion that interactions between the hippocampus and default network regions are important for long-term encoding and retrieval of semantically rich real-world events.

Within the default network regions of D.A.’s brain, activity patterns in PPC and PMC during narrative listening were relatively preserved. In contrast, activity patterns in mPFC were less preserved. PPC and PMC also demonstrated the “unscrambling” effect in the amnesic brain, while mPFC did not. Interestingly, among default network regions, mPFC is known to be strongly coupled with the hippocampus in studies of resting state connectivity (Kahn et al., 2008) and oscillatory synchrony; in particular, hippocampal-PFC interactions are thought to be crucial for learning and remembering events (Eichenbaum, 2017; van Kesteren et al., 2012). Thus, it may be that the missing interactions with hippocampus caused mPFC activity to diverge from the healthy pattern more than PPC or PMC. It should be noted that negative results are difficult to interpret, as differences between the amnesic and healthy brain could arise from a number of plausible sources, such as undiagnosed atrophy or the substantial age difference between the patient (63) and the controls (18–31). At the same time, these plausible sources of differences make positive results, where significant similarities between patient and controls are observed, more challenging to attain. D. A.’s default network brain activity (PPC and PMC, but not MPFC) during narrative listening was significantly similar to a healthy control cohort over three decades younger than him, in spite of his dense amnesia.

Our observations may also have implications for the phenomenological experience of amnesics when they listen to stories or watch movies that are multiple minutes long: although they are later unable to report the details, their moment-by-moment comprehension of narratives may be similar to that of healthy individuals in many respects. It has been challenging to concretely assess amnesic patients’ experience of minutes-long continuous stimuli like stories. Prose tests tend to be very short, typically less than one minute; the Logical Memory Test component the Wechsler Memory Scale (Wechsler, 1987), a widely-used standard measure of prose memory, is only 65 words long. Amnesic immediate recall performance for short prose passages, while not as catastrophically impaired as after a filled delay, is still poor (Baddeley and Wilson, 2002; Rosenbaum et al., 2008). In verbal discourse about future and past events, amnesics show impairments in both low and high level coherence (Kurczek and Duff, 2011; Race et al., 2015a). One challenge in conducting such behavioral tests is that interrupting story-listening to ask questions changes the context and necessarily creates a discontinuity; in other words, testing recall can introduce a discontinuity which hinders recall (Jang and Huber, 2008). Thus, while suggestive, extant behavioral studies do not show that amnesics can follow a narrative multiple minutes long. Our finding that a 63-year-old hippocampal amnesic’s brain activity patterns resemble those of young (18–33) healthy controls, moment-by-moment over the course of a 7-minute narrative, suggests that recall tests have underestimated the ability of hippocampal amnesics to integrate naturalistic information over time.

In this paper we present evidence that the purported ability of default network cortical areas to integrate information across long timescales, on the order of 30 seconds or more, does not depend solely on interactions with the hippocampus. We scanned an amnesic patient with bilateral hippocampal damage as he listened to intact and paragraph-scrambled auditory narratives. Within the default network (PPC and PMC, but not MPFC), the amnesic’s moment-by-moment brain activity patterns were significantly similar to that of a healthy control cohort; in the same brain areas, responses were affected by the story scrambling, indicating that they integrated information across paragraphs. The observations in this case study support the idea that default network cortical areas have an intrinsic ability to carry slowly-changing information during semantically rich natural input.

## Acknowledgements

We thank Lok-Kin Yeung and Rachel Newsome for assistance with data collection; we thank Buddhika Bellana, Hongmi Lee, and Lisa Musz for their comments on the manuscript. JC was supported by the Sloan Research Fellowship. CH was supported by the Natural Sciences and Engineering Research Council of Canada (RGPIN-2014-04465) and the Sloan Research Fellowship. MB was supported by the James S. McDonnell Scholar Award. KN and UH were supported by the National Institute of Mental Health award R01MH112357.

