## Supplementary Materials for "Temporal integration of narrative information in a hippocampal amnesic patient"

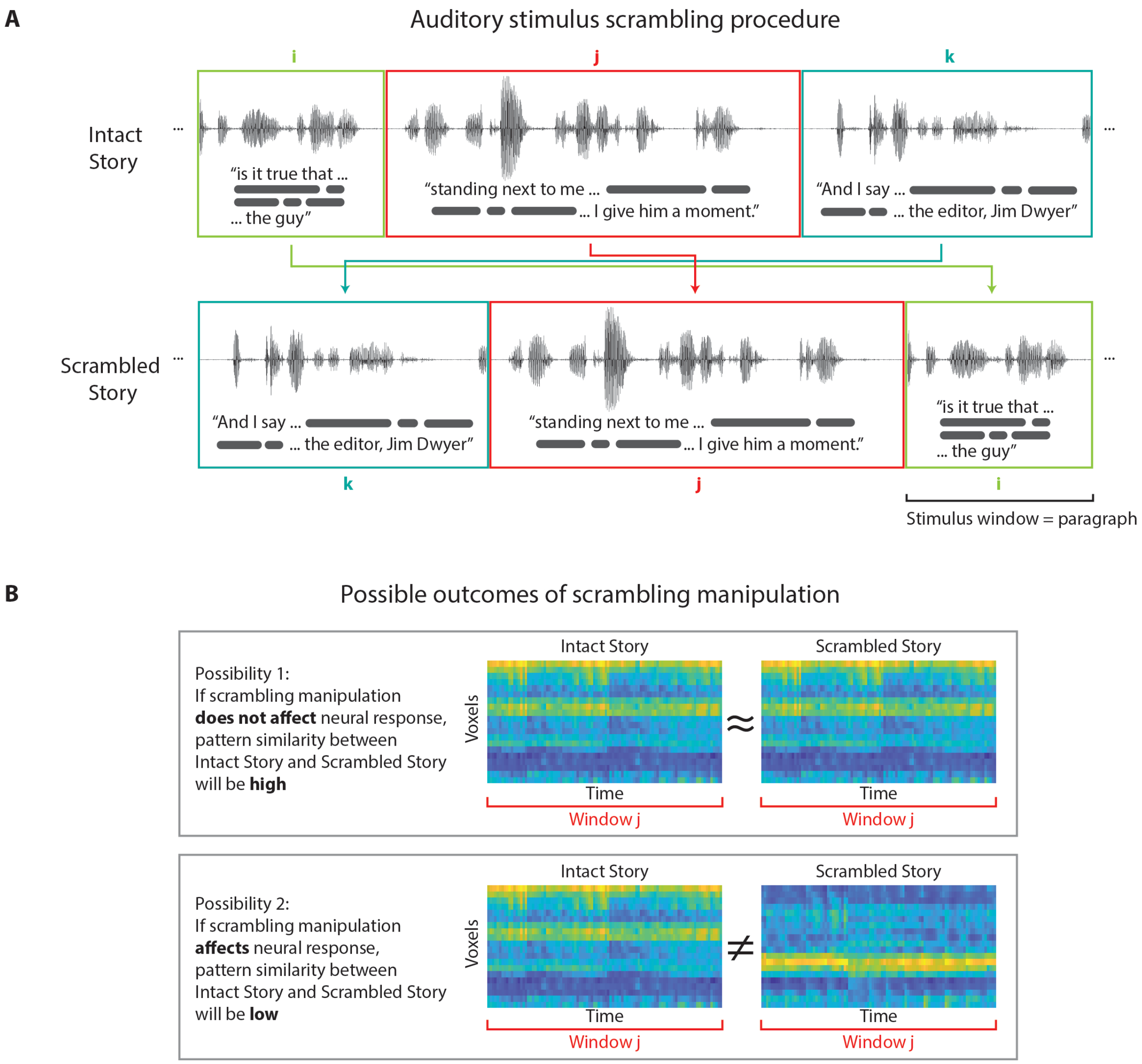

**Supplementary Figure 1.** Scrambling schematics. **A**) Schematic showing the scrambling procedure for auditory stimulus. **B**) Schematic showing the pattern match of corresponding stimulus windows between the Intact Story and Scrambled Story.

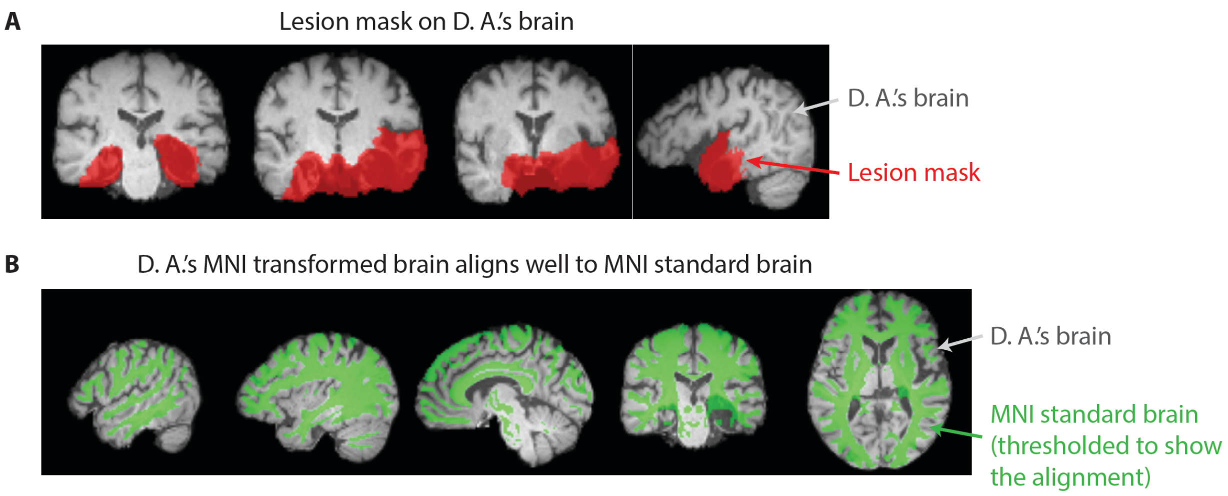

**Supplementary Figure 2.** Patient anatomy and lesion mask. **A**) Experimenter defined lesion mask (red) on D. A.'s anatomical image (grey) in standard 2-mm MNI space. **B**) 2-mm MNI brain (green; intensity threshold at 6500–8000) on D. A.'s anatomical image (grey) to show the match of the transformation.

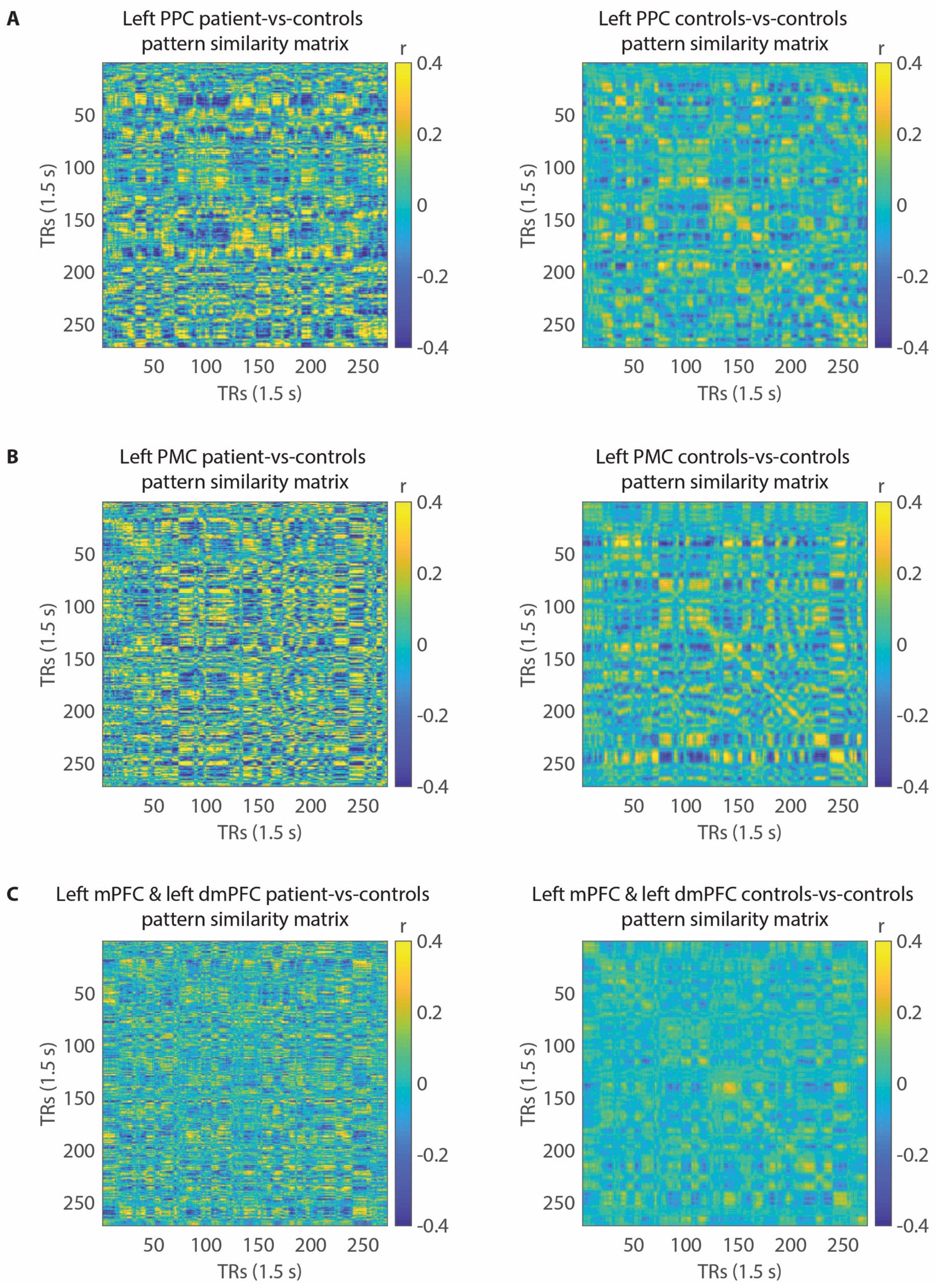

A
Left PPC patient-vs-controls
pattern similarity matrix
Left PPC controls-vs-controls
pattern similarity matrix
B
Left PMC patient-vs-controls
pattern similarity matrix
Left PMC controls-vs-controls
pattern similarity matrix
C
Left mPFC & left dmPFC patient-vs-controls
pattern similarity matrix
Left mPFC & left dmPFC controls-vs-controls
pattern similarity matrix
r
0.4
0.2
0
-0.2
-0.4
TRs (1.5 s)
50
100
150
200
250

**Supplementary Figure 3.** Pattern similarity matrices in selected ROIs between the patient and controls as well as within control subjects. **A**) Left posterior lateral parietal cortex (PPC). **B**) Left posterior medial cortex (PMC). **C)** Combined left medial prefrontal cortex (mPFC) and dorsal medial prefrontal cortex (dmPFC). Values in the diagonals of the matrices (top left corner to bottom right corner) are the true pattern correlation values calculated between aligned time points. Off-diagonals indicate pattern correlation values between time points that are not aligned. Controls-vs-controls pattern similarity is highest along the diagonals, suggesting that the pattern match among control subjects was temporally specific, i.e., dependent on the stimulus matching moment-by-moment.

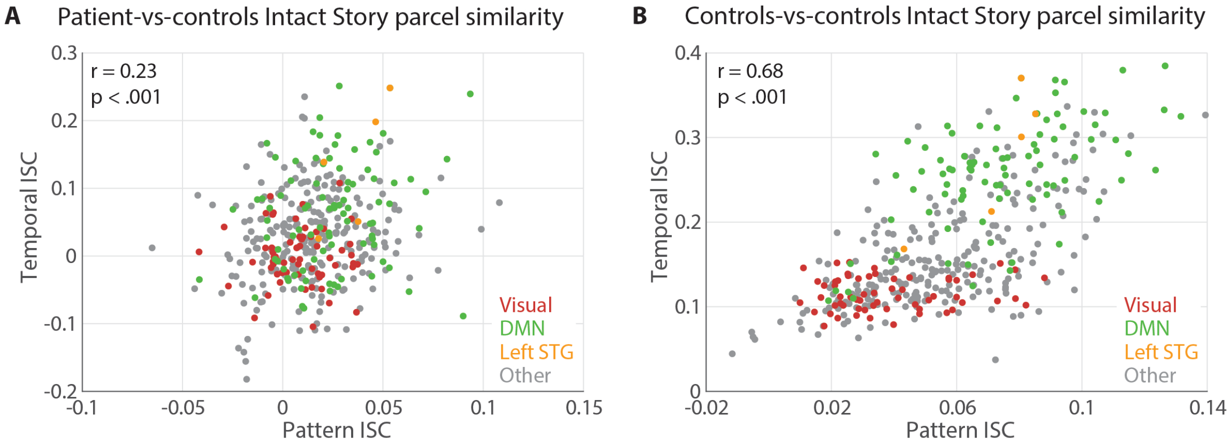

**Supplementary Figure 4.** Intact Story pattern and temporal similarity by parcels. **A**) Patient-vs-control pISC and tISC are positively correlated ($r = 0.23$, $p = 4.8 \times 10^{-6}$). **B**) Controls-vs-controls pISC and tISC are positively correlated ($r = 0.68$, $p = 4.8 \times 10^{-55}$). Visual network parcels have relatively lower pISC and tISC (due to the stimulus being non-visual), whereas auditory and default network parcels have relatively higher ISC.

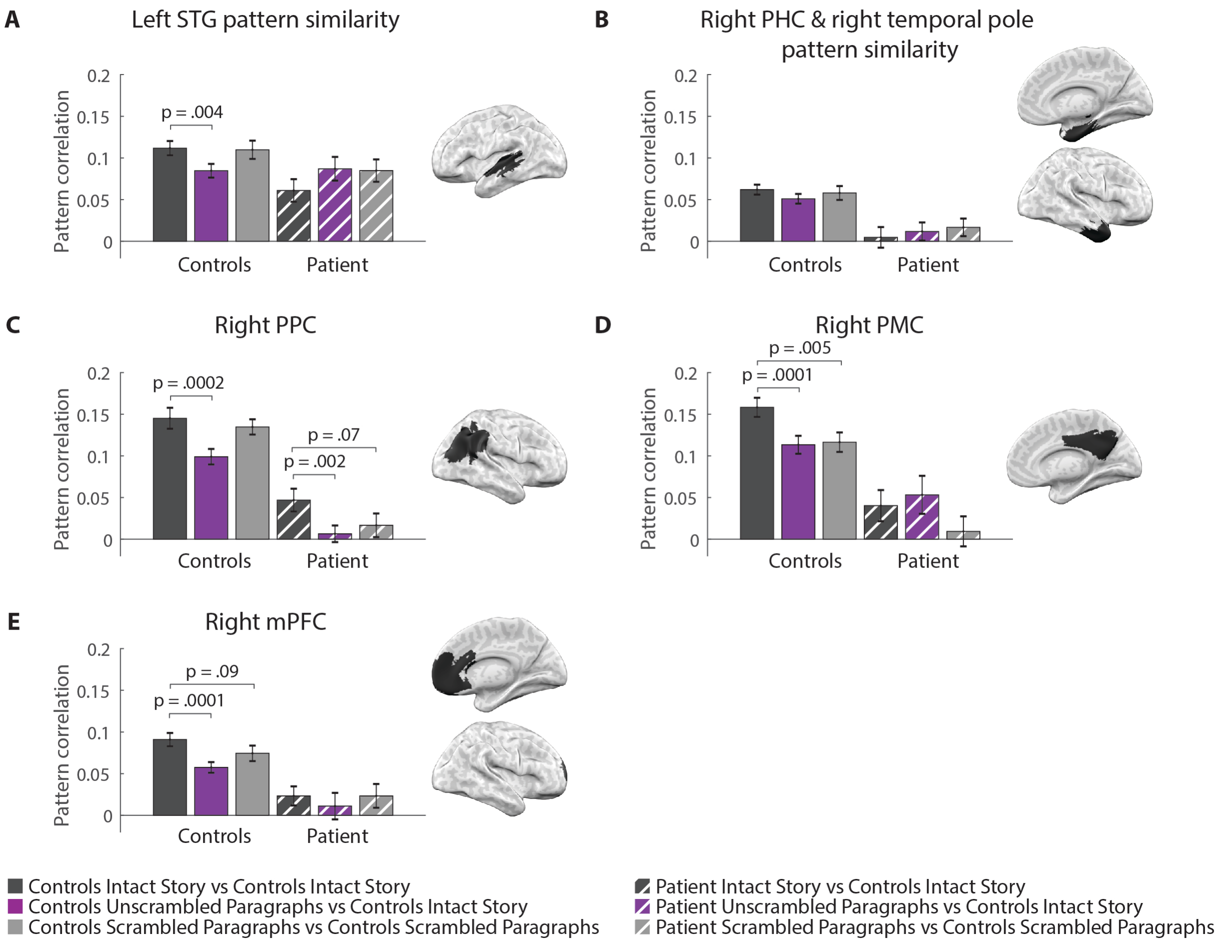

**Supplementary Figure 5.** Pattern similarity in selected ROIs. **A**) Left superior temporal gyrus (STG). **B**) Combined right parahippocampal cortex (PHC) and temporal pole ROIs. Error bars indicate bootstrapped standard error. Bootstrapped p values are labeled in the figure if $p < 0.1$.

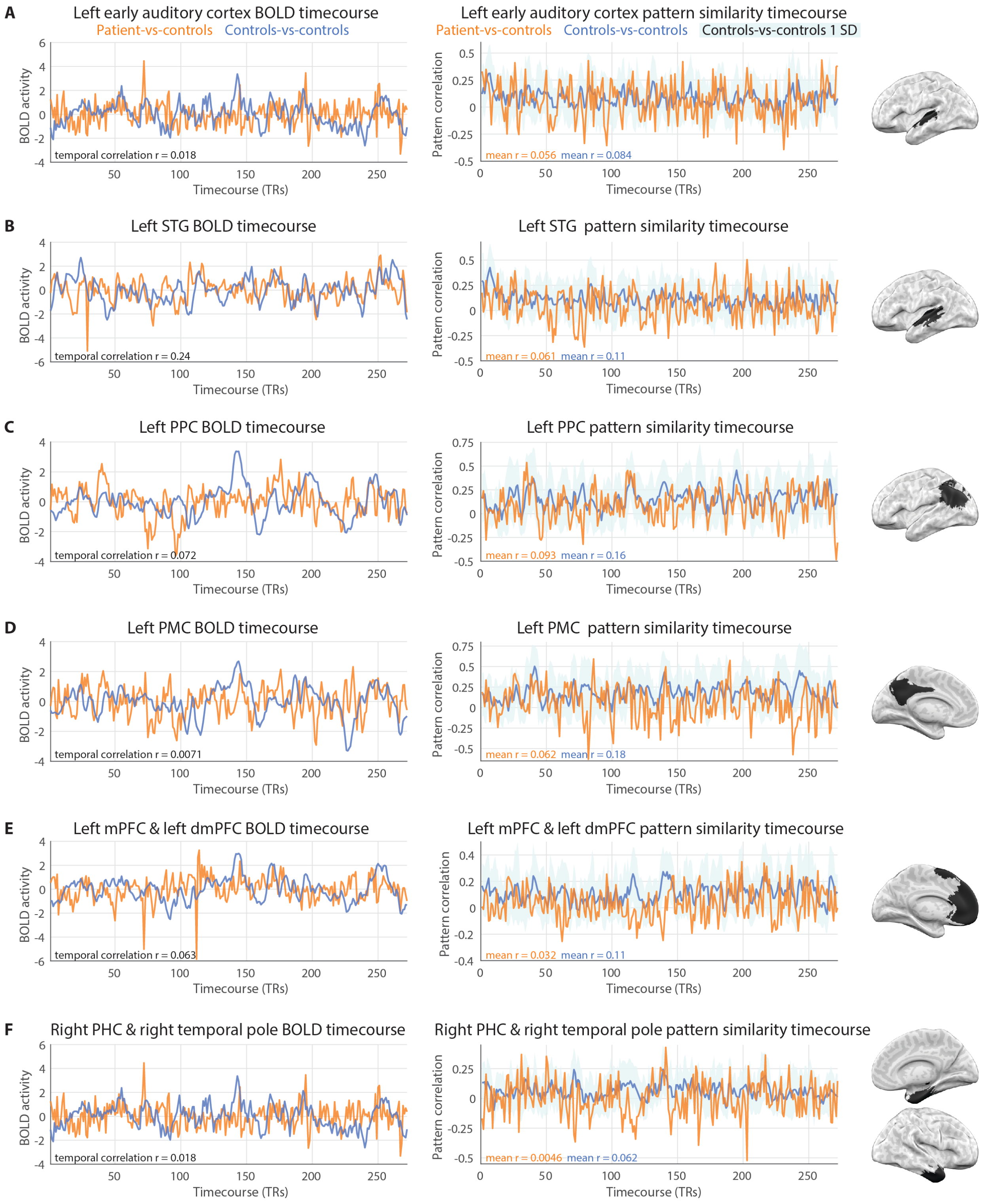
A
Left early auditory cortex BOLD timecourse
Patient-vs-controls
Controls-vs-controls
BOLD activity
temporal correlation r = 0.018
Timecourse (TRs)
Left early auditory cortex pattern similarity timecourse
Patient-vs-controls
Controls-vs-controls
Controls-vs-controls 1 SD
Pattern correlation
mean r = 0.056
mean r = 0.084
Timecourse (TRs)
B
Left STG BOLD timecourse
BOLD activity
temporal correlation r = 0.24
Timecourse (TRs)
Left STG pattern similarity timecourse
Pattern correlation
mean r = 0.061
mean r = 0.11
Timecourse (TRs)
C
Left PPC BOLD timecourse
BOLD activity
temporal correlation r = 0.072
Timecourse (TRs)
Left PPC pattern similarity timecourse
Pattern correlation
mean r = 0.093
mean r = 0.16
Timecourse (TRs)
D
Left PMC BOLD timecourse
BOLD activity
temporal correlation r = 0.0071
Timecourse (TRs)
Left PMC pattern similarity timecourse
Pattern correlation
mean r = 0.062
mean r = 0.18
Timecourse (TRs)
E
Left mPFC & left dmPFC BOLD timecourse
BOLD activity
temporal correlation r = 0.063
Timecourse (TRs)
Left mPFC & left dmPFC pattern similarity timecourse
Pattern correlation
mean r = 0.032
mean r = 0.11
Timecourse (TRs)
F
Right PHC & right temporal pole BOLD timecourse
BOLD activity
temporal correlation r = 0.018
Timecourse (TRs)
Right PHC & right temporal pole pattern similarity timecourse
Pattern correlation
mean r = 0.0046
mean r = 0.062
Timecourse (TRs)

**Supplementary Figure 6.** BOLD activity timecourse and pattern similarity timecourse in selected ROIs. **A**) Left early auditory cortex. **B**) Left superior temporal gyrus (STG). **C**) Left posterior lateral parietal cortex (PPC). **D**) Left posterior medial cortex (PMC). **E**) Combined left medial prefrontal cortex (mPFC) and dorsal medial prefrontal cortex (dmPFC). **F**) Combined right parahippocampal cortex (PHC) and temporal pole ROIs.

**Supplementary Table 1. List of stories used as auditory stimuli**

| Experiment | Story | Author/Narrator | Producer | Link |
|---|---|---|---|---|
| fMRI | Pieman | Jim O'Grady | The Moth | https://themoth.org/stories/pie-man |
| Long Story Segments | Smalltalk | Kevin Carlin | The Moth | https://themoth.org/stories/smalltalk |
| Long Story Segments | You are a Great King | Alex Draper | The Moth | https://themoth.org/stories/you-are-a-great-king |
| Short Stories | Travels With My Cats | Written by Mike Resnick, read by Serah Eley | Escape Pod (EP082) | http://escapepod.org/2006/11/30/ep082-travels-with-my-cats/ |
| Short Stories | The Ludes | Written by Lisa M. Bradley, read by Serah Eley | Escape Pod (EP026) | http://escapepod.org/2005/11/03/ep026-the-ludes/ |
| Short Stories | Stu | Written by Bruce McAllister, read by Serah Eley | Escape Pod (EP135) | http://escapepod.org/2007/12/06/ep135-stu/ |
| Short Stories | Article of Faith | Written by Mike Resnick, read by Serah Eley | Escape Pod (EP193) | http://escapepod.org/2009/04/02/ep193-article-of-faith/ |
| Short Stories | Revolution Time | Written by Lavie Tidhar, read by Serah Eley | Escape Pod (EP163) | http://escapepod.org/2008/06/20/ep163-revolution-time/ |
| Short Stories | The Shoulders of Giants | Written by Robert J. Sawyer, read by Serah Eley | Escape Pod (EP078) | http://escapepod.org/2006/11/02/ep078-the-shoulders-of-giants/ |
| Short Stories | Hell Notes | Written by M.K. Hobson, read by Serah Eley | Escape Pod (EP015) | http://escapepod.org/2005/08/18/ep015-hellnotes/ |
| Short Stories | Mr. Penumbra's Twenty-Four-Hour Book Store | Written by Robin Sloan, read by Serah Eley | Escape Pod (EP215) | http://escapepod.org/2009/09/10/ep215-mr-penumbras-twenty-four-hour-book-store/ |
| Short Stories | Love and Death in the Time of Monsters | Written by Frank Wu, read by Serah Eley | Escape Pod (EP167) | http://escapepod.org/2008/07/18/ep167-love-and-death-in-the-time-of-monsters/ |
| Short Stories | Blood of Virgins | Written by David Barr Kirtley, read by Serah Eley | Escape Pod (EP088) | http://escapepod.org/2007/01/11/ep088-blood-of-virgins/ |
| Short Stories | End Game | Written by Nancy Kress, read by Serah Eley | Escape Pod (EP125) | http://escapepod.org/2007/09/27/ep125-end-game/ |

**Supplementary Table 2. Behavioral experiment listen and recall instructions**

| Session | Instructions |
|---|---|
| Long Story Segments | "In this section you will be listening to stories broken up into segments, and repeating what you heard. Your response does not have to be word-for-word, but should include as much detail as possible." |
| Sentence Pairs | "In this section you will be listening to pairs of sentences, and repeating them word-for word." |
| Short Stories | "In this section you will be listening to short narratives, and repeating what you heard. Your response does not have to be word-for-word, but should include as much detail as possible." |
| WMS | "In this section you will be listening to short paragraphs, and repeating them word-for word." |

**Supplementary Table 3. List of Schaefer 400 parcels used to create region of interests**

| Region of interest | Schaefer parcel ID | Notes |
|---|---|---|
| Left early auditory cortex | 44 45 46 | Parcels closest to primary auditory cortex in “Somato-Motor” network |
| Left superior temporal gyrus (STG) | 44 45 46 196 197 | Auditory cortex and two adjacent parcels in “Temporal-Parietal” network |
| Left posterior lateral parietal cortex (PPC) | 149 150 173 174 188 136 137 138 | All parcels labeled “IPL” in the three “Default” networks; “IPL” in “ContB” network |
| Left posterior medial cortex (PMC) | 154 155 156 157 158 159 160 | All parcels labeled “PCC” in “DefaultA” network |
| Combined left medial prefrontal cortex (mPFC) and dorsal medial prefrontal cortex (dmPFC) | 161 162 163 164 165 166 175 176 177 178 179 180 | All parcels labeled “PFCm” in “DefaultA” network; all parcels labeled “PFCd” in “DefaultB” network |
| Combined right parahippocampal cortex (PHC) and temporal pole ROIs | 389 319 320 322 376 | “PHC1” in “DefaultC” network; “TempPole” 1, 2 & 4 in “Limbic” network; “AntTemp” in “DefaultB” network |
| Right posterior lateral parietal cortex (PPC) | 359 360 385 386 338 339 340 341 304 | Parcels labeled “IPL” in “DefaultA”, “DefaultC”, “SalVentAttnB”, and “ContB” networks |
| Right posterior medial cortex (PMC) | 363 364 365 366 367 | All parcels labeled “PCC” in “DefaultA” network |
| Right medial prefrontal cortex (mPFC) | 368 369 370 371 372 373 | All parcels labeled “PFCm” in “DefaultA" network |

**Supplementary Note 1: Auditory stimuli transcripts**

**1. Long Stories transcript**

Story 1: You are a Great King

Segment 1

Uh I was doing a play at the Olney Theater, Olney Maryland, and one down to break from rehearsal, a woman from the developmental office poked her head in the room and said, “Alex, it’s India, on Line 1 for you.” I went to the phone and it was my friend Govind Menon. Govind is an Indian friend who I went to college with and did many plays with in college, and he was calling me from India to tell that he was cast in but also helping to produce a huge Indian movie, and he was calling me up to see if I was available for two months to come to India and be in the movie.

Segment 2

He was being very vague and very pushy, uh it was a huge movie but there was no budget, uh. There were huge stars in it, well like who? You wouldn’t know them. Uh, can I read a script? There’s no script. Throughout all his vagueness, the one hook that he had was the role he wanted me to play. He told me that it was a prison movie and he wanted me to play the warden. And this warden was a man who actually existed, a real-life person who existed.

Segment 3

And who, as a matter of historical fact, had the following very tantalizing to an actor, character attributes: uh he was Irish, lovely; he had a horrible limp, and used a cane, fantastic; he was hard of hearing and used an ear trumpet, colorful; he was psychopathically violent, oh come on; he was an alcoholic, fantastic; and he was really, really fat.

Segment 4

And I told Govind I said, “Govind, I’m your man, but I’m not fat.” “Are you a little bit fat?” “No I’m not fat at all.” “Are you like, chubby?” “Govind I’m not fat.” “Can you get fat?” “Govind I can’t get fat! You won’t even send me a script, I can’t get fat!” “Look, don’t worry about it. You’re gonna be great. They’ll love you.” So after I flew to India, about half way through the 24-hour flight, I started having serious doubts about what I’ve decided to do. I had no idea what I was getting myself into. It was, It was two months, I didn’t know where I was going; I didn’t know any of the people I was going to be working with; I had no idea what this movie was about.

Segment 5

But through all that I figured you know what, if nothing else, I’m going to the Andaman and Nicobar Islands in the Bay of Bengal, South of Burma, if nothing else it’ll be an adventure. For the last leg of the trip, from Madras to Port Blair, Govind and one of the movie stars and two of the producers joined me on the plane. And when we landed in Port Blair, this tiny little one-strip airstrip, uh there was a huge crowd waiting for us. And the plane pulled to a stop and the crowd surrounded the plane and they, you know, they couldn’t get the little stairs through and they finally did.

Segment 6

I'm looking out the window when I see five guys in baseball caps and sunglasses dart down the stairs and they, they got ushered into this van that's waiting there, and they sort of pulled away and the crowd goes nuts and follows them and…and I turned to Govind and I say, "Govind, who are those guys?" And he kind of smiles and said, "that's us man." Like, "what do you mean?" "That's us, decoys. They're pulling the crowd away from the plane so that we can get off."

Segment 7

And again I think that maybe this thing was bigger than I thought. That night, there was a party in the director's suite, at this—we're in this beautiful hotel on this beautiful island and there's a party that night in the director's suite, and it's a—it's a party because that day, the winners of the Indian Academy Awards were announced. And people connected to the movie had won 17 academy awards. So I was getting very nervous, and I had to go and meet the director. And we go meet him and it's a room full of large Indian men in really nice shirts and Rolexes and Ray-Bans and…

Segment 8

...and they all seem very nice. They don't speak much English but they're sort of quiet and focused and nice. I thought it went well, and later, in my hotel room, I told Govind that I thought it went well and he said, "nah that didn't go well at all." So I, "what do you mean?" "They're really upset, man." "What're they…" "They're very upset that you're not fat."

Story 2: Smalltalk

Segment 1

Um, south of Buffalo, New York, there is a ravine that's called Zoar Valley. And if you park, and hike a mile up the creek, there is a wat—there's a twenty-foot waterfall that you can jump off of, and it is amazing. And as you can imagine, a lot of people find that jumping off a waterfall is a little too boring and mundane unless they involve alcohol. And so, not surprisingly, each summer there are an unfortunate number of injuries, including the occasional death. And, the last time I was there with my friend Justin, we had just jumped the falls, we were walking back to the car. And around the first bend we came across a drunk college kid who had tried to climb up the ravine, and it's just loose shale so of course he fell and, like, hit the back of his head. And, uh, by the grace of god, Justin and I were the second people to come across this kid. Had we been the first people to find him, we almost certainly would have just watched him expire, 'cause we don't know what we're doing. But, uh, by some miracle the–

Segment 2

But, uh, by some miracle the first guy to find this kid was an army field medic, so he knew exactly what he was doing. So he was in his element. And he had the kid- He had him lying on his back, on the rocks, with a towel over him. And, uh, as we came up, he was telling him, like: "look, you got a scratch on the back of your head, we gotta keep you stabilized, we gotta keep you warm. You're gonna be fine, and we're gonna get you help." And, uh, so we walk up and the guy's, like: "we gotta keep this kid warm." So we took my towel, and we got it under him to insulate him from the rocks beneath him, and then Justin's towel we rolled up and put behind his head. And then the medic takes me aside, and he goes: "this kid has a gigantic gash on the back of his head." And he goes "I gotta look at it. What I need you to do is I need you to keep him focused, and keep him talking, because the instant he closes his eyes, he's going to die."

Segment 3

Never did I think that I would be in a crisis situation, where the life-or-death task I would be given, would be to make small talk with a drunk college kid. Didn't think it would had happened. I was not prepared for that, I am an absolute introvert. Small talk is not my forte. This kid lives, but you should, all of you, nevertheless pray that your life never depends on my ability to have a skin-deep conversation. They are long odds. And so I, uh, I dug deep, I made the only kind of small-talk I ever knew how to make, which is very awkward. It was to ask him the sort of questions I would ask a girl I was interested in getting to know better.

Segment 4

And I haven't even had to do that in a long time, 'cause I'm married. So I was—I had to dig deep. It was awkward, I was like: "So are you in school? Where do you go to school?" And he's like: "UB." And I was like: "Oh, UB, that's great man. Are you majoring? What's your- What are you studying?" And he's like- he's like: "History. I'm so cold man, I'm really cold." And then like, all I could think to do was just ignore that. So I was like: "Oh, history, that's interesting, that's- What do you think you wanna do when you graduate? You wanna teach? You wanna be a teacher?" He's just like: "I'm just really cold man." I was just like: "Oh, we should get together, we should have a history study session some time." And, like, he's, like, literally like starting to do the, like, eyes-half-closed thing, and I was just like in an absolute panic. So it started to deteriorate for me."

Segment 5

And I started to just, like, get mad. And I was just li- It was like a police interrogation at that point. I was just like: "I asked you a question, son!" Trying anything. And then in the meantime, while this was going on, Justin had been given the task of finding a cell-phone signal and calling 911. And I was more than a little envious of that job. Um, I happen to be very good at that. You need a- you need a cell-phone signal in a ravine, I'm your guy: cell-phone-signal-in-a-ravine-finding-fool. In fact my ego was a little bruised that I wasn't selected for that. But, uh, he got a hold of 'em, and so they dispatched a helicopter to air-lift the kid out. And, um, in the meantime, while the helicopter was en route, two for- er, uh, park rangers hiked up to us with medical supplies, and they kinda took over.

### 2. Sentence Pairs transcript

#### Coherent sentence pairs

| Sentence Pair Number | Sentence A | Sentence B |
|---|---|---|
| 1 | The professor was finishing handing out the exams. | The examination room began to fill with the sound of scratching pencils. |
| 2 | Emily scowled when she saw who had entered the weight room. | The rivals had competed fiercely on the track many times. |
| 3 | Carroll knew the kids would love the cake she had baked. | The children sprinted to the table as the candles were being lit. |
| 4 | Tony led his son to their seats behind home plate. | With peanuts and cotton candy, this was going to be a memorable game. |
| 5 | The librarian's cart rattled loudly as it passed by. | Jim woke up suddenly with his geology textbook in his hands. |

| | | |
|---|---|---|
| 6 | The tour began with a cheesy video of the soda factory's history. | Kevin was beginning to think that the outing wasn't worth the expense. |
| 7 | Patricia and her friends from work decided not to get a golf cart. | The course was a long eighteen holes, but the exercise couldn't hurt. |
| 8 | The mechanics worked hard, stopping only to wipe grease from their faces. | Everybody was anxious to hear the antique car roar to life. |
| 9 | The riot police issued commands over the loudspeakers. | Protesters began to leave the streets as backup officers arrived. |
| 10 | Suzan caught a glimpse of the beach through the airplane window. | Exotic coconut trees were growing all along the coastline. |
| 11 | The museum curator was proud to announce the new exhibit. | The rare Aztec artifacts would draw an enthusiastic crowd. |
| 12 | Doctor Wilson leaned over and carefully read the clipboard. | The patient's appendix removal surgery had gone well. |

**Incoherent sentence pairs**

| Sentence Pair Number | Sentence A | Sentence B |
|---|---|---|
| 13 | Heavy waves sprayed over the bow of the sailboat all day. | As the subway arrived, the wind blew the hat off a young girl. |
| 14 | Lisa stood in the goal, waiting for the penalty kick. | The poodle sprinted out the door before the leash could be attached. |
| 15 | The explorers felt pride as they planted their flag in the Arctic ice. | The movie previews had been lengthy, but the feature was about to begin. |
| 16 | Rosa was bumped many times as she moved along the busy street. | A lone eagle soared above the crevasse of the Colorado river. |
| 17 | The group had been ascending the trail for six hours. | The jugglers had been great and the trapeze was next up. |
| 18 | He walked through the cafeteria trying to choose something for dinner. | The passengers smiled childishly as the roller coaster came to a halt. |
| 19 | The audience waited as the pianist settled her hands on the keys. | Jerry flicked through the channels finding nothing but commercials. |
| 20 | The robot crawled over the surface of the barren planet. | The detective was completely baffled by the empty bank vault. |
| 21 | The minister greeted everyone as they entered for Sunday service. | Doctor Chang took a sample from his expedition back to the lab. |
| 22 | The field guide hacked through the thick jungle vines. | Dinner had been nice, but the restaurant's prices were ridiculous. |
| 23 | The sun set behind the pyramids as they finished their picnic. | Everyone tidied their desks on the day that the president visited. |
| 24 | The notorious criminal faced the jury without regret. | Inside the covered market the stench of fish was unbearable. |

**Sentence word count**

| Sentence Pair Number | Sentence A Word count | Sentence B Word count |
|---|---|---|
| 1 | 8 | 12 |
| 2 | 11 | 10 |
| 3 | 11 | 12 |
| 4 | 10 | 13 |
| 5 | 9 | 11 |
| 6 | 12 | 12 |
| 7 | 13 | 12 |
| 8 | 12 | 11 |
| 9 | 8 | 10 |
| 10 | 11 | 9 |
| 11 | 10 | 9 |
| 12 | 9 | 8 |
| 13 | 11 | 13 |
| 14 | 10 | 12 |
| 15 | 13 | 13 |
| 16 | 12 | 11 |
| 17 | 10 | 11 |
| 18 | 11 | 12 |
| 19 | 12 | 9 |
| 20 | 10 | 10 |
| 21 | 10 | 12 |
| 22 | 9 | 10 |
| 23 | 11 | 11 |
| 24 | 8 | 10 |

**3. Short Stories transcript**

Travels With My Cats (Congruent)

That night I was faced with a major decision. I didn't want to read a book called "Travels with my Cats" by a woman called "Miss", but I'd spent my last nickel on it—well, the last until my allowance came due again next week—and I'd read all my other books so often you could almost see the eye tracks all over them. So, I picked it up without much enthusiasm and read the first page, and then the next, and suddenly I was transported to Kenya Colony, and Siam, and the Amazon. Miss Priscilla Wallace had a way of describing things that made me wish I was there, and when I finished a section, I felt like I'd been there. There were cities I'd never heard of before, cities with exotic names like Maracaibo and Samarkand and Addis Ababa. Some with names like Constantinople that I couldn't even find on the map. Her father had been an explorer. Back in the days when there still were explorers. She had taken her first few trips abroad with him, and he had undoubtedly given her the taste for distant lands. My own father was a typesetter. How I envied her. I had half hoped the African section would be filled with rampaging elephants and man eating lions, and maybe it was, but that wasn't the way she saw it. Africa may have been red of tooth and claw, but to her it reflected the gold of the morning sun, and the dark shadowy places were filled with wonder, not terror. She could find beauty anywhere. She would describe 200 flower sellers lined up along the Seine on a Sunday morning in Paris. Or a single, frail blossom in the middle of the Gobi Desert, and somehow you knew that each was as wondrous as she said. And suddenly I jumped as the alarm clock started buzzing. It was the first time I'd ever stayed up for the entire night. I put the book away, got dressed for school, and hurried home after school so I could finish it. I must have read it six or seven more times that year. I got to the point where I could almost recite parts of it word for word. I was in love with those exotic far-away places, and maybe a little bit in love with the author too. I even wrote her a fan letter addressed to "Miss Priscilla Wallace, Somewhere", but of course it came back.

The Ludes (Congruent)

Back then, my favorite thing to do was to pop a few tranquilizers and sit in the balcony for student recitals. As long as I sat up there, out of sight, no one seemed to care if I drowsed and drooled the entire time, or slumped in my licorice-red seat and stared at the ceiling in a pharmacological daze. And, as these little concerts were free, it seemed as good a way as any to pass the cold winter nights in a boring little college town. One night as I slouched there, waiting for the music to begin, I noticed a young man enter the balcony. I noticed him, despite the drug-induced haze gently drifting over me, because I'd never seen him before, and he didn't belong. That might seem funny to those who've ever bothered to attend these performances, to say that someone didn't belong. The audience is always a motley sort: faculty and spouses, local musicians and artists, waitresses and office workers desperate for some culture, and of course, recreational drug users with nothing better to do. Still, he didn't belong. He was Gothic. I'd seen goths there before, and he wasn't goth, he was Gothic. Dark and looming, faintly chivalrous in manner, seemingly possessed of a great tragic secret. Then the cellist appeared onstage, and after a predatory moment of silence, he sank into the prelude. My haze turned into a dense fog; a warm gooey feeling came over me. Between the prelude and allemande were the usual fidgetings of the audience. A muffled cough here, someone shrugging out of a sweater there. Then, out of the corner of one drooping eye, I noticed movement in the balcony. The Gothic man was leaving, a blue bandana pressed to his face, as if he were weeping. I was impressed. To be so moved by the music, when I merely luxuriated in it, like a pig in a blanket.

Stu (Congruent)

Stu lived up near Escondido, in the avocado orchards, and it took 45 minutes. When we got there, he had this gigantic plastic above-ground swimming pool in his backyard, and this machine suspended above it on a winch, the kind you use to lift engines out of cars. No one said a thing about the pool as we ate hotdogs, hamburgers, and potato salad in the backyard. But in the car driving home, our dad said, "He's using it to look for oil." "What?" we both said. He looked at us slyly and said, "He's using sonar. Sonar aimed down through the water in the pool, to look for oil. No one's ever done it before. The Navy's not interested in using it that way, which is why he can talk about it. But let's still keep it our little secret, okay boys?" We always kept things secret. When you're not even supposed to carry one of your dad's "Property of US Government" ballpoint pens to school, you learn to keep secrets—even ones the Navy isn't interested in. "Sure, dad!" we said. "If it works, won't he get rich?" I asked. I was the oldest, so I asked questions like that. "No, he works for the Navy. Anything he invents belongs to them, whether they're interested in it or not." The next time we visited Stu, I was twelve, and the big plastic above-ground pool was gone. So was the machine with its winch. My brother and I looked around the yard for anything that looked like an invention, and couldn't see a thing. But, when we went inside for dinner—Stu loved to barbecue, so it was patties and wieners again—there was Stu, standing in the middle of the living room, holding something, and grinning. His white hair stuck out like Einstein's, and though he didn't wear a moth-eaten sweater like Einstein, he wore this little vest that looked just as sill y. Stu's face was a little crazier looking too, his eyes were open a little too wide, as if something had just bitten him, and his smile was a little crooked. He was holding a machine about the size of a workman's lunch pail, and when we saw how he was grinning at us, we knew he'd waited for us to come in before telling our mom and dad about it.

Article of Faith (Congruent)

He went off to get the water, and I went to my office, hung my coat up in the closet, opened the center drawer of my desk, and pulled out my sermon. I have a wonderful, state-of-the-art computer, that can probably think a thousand times faster than I can. But somehow, I'm more comfortable writing out my sermons longhand, on a legal pad. I made a couple of last-minute changes, and left the office. A minute later, I was standing in the pulpit, clutching the podium with both hands as I always do. If I don't, I tend to gesticulate too much. And I began working my way through the sermon. When I finished, I checked my watch. It had taken 22 minutes, which seemed an acceptable length. My rule of thumb has always been that anything over 30 minutes is likely to be boring, and anything under 15 minutes seems truncated and insufficiently thoughtful. I looked up from my watch and saw Jackson standing motionless at the back of the church. "I'll get out of your way now," I said, starting to walk back toward my office. "Continue whatever you were doing." "Yes, Reverend Morris," said Jackson. Then a thought occurred to me. "Just a minute, Jackson." "Sir?" "Were you able to hear my sermon?" "Yes, Reverend Morris." I looked at him. "Well, what did you think of it?" "I do not understand the question." "Let me explain then," I said. "I give a sermon to my parishioners every Sunday morning. It is supposed to bring them spiritual comfort. But it is also intended to instruct them." "Instruct them, sir?" said Jackson. "On how to lead moral, spiritually satisfying lives," I explained. "The problem is that sometimes I get too close to my subject matter, and I don't see any logical flaws or contradictions that have crept in." I smiled at him. "I would like you to listen to my sermons, not on Sunday mornings, but when I am practicing them during the week, and point out any logical inconsistencies in them. Do you think you can do that?"

Revolution Time (59 s) / The Shoulders of Giants (55 s) (Incongruent)

So I did my part. Photographed the building, the outlying streets, possible entrances, how often the guards changed. Grunt work, and redundant besides, since everyone knows the real body of the Institute is several miles away, where the town ends and the desert begins. Like Area 51 in its time, it's a badly kept secret. I'd already spent over a week sidetracked to this part of the operation. It wasn't only the building of course, I made note of the researchers and politicians who came through the building, making detailed photographic records of each one. I knocked on doors in various pretenses, to try and determine possible observation posts, escape routes, pick up rumors and gossip which may prove useful. Satisfied at last that I could finish my watch, I made my way with the other tourists to the subway station, and took the train to the cell's meeting place. There were four members in my cell. I was the only local, but it was Joe who was going to get us in.

My father had been from Glasgow, my mother from Los Angeles. They had both enjoyed the quip that the difference between an American and a European was that to an American, a hundred years was a long time, and to a European, a hundred miles is a long journey. But both would agree that 12 hundred years and 11.9 light years were equally staggering values. And now here we were, decelerating in toward Tau Ceti, the closest sun-like star to Earth that wasn't part of a multiple star system. Of course, because of that, this star had been frequently by Earth's search for extra-terrestrial intelligence. But nothing had ever been detected. Nary a peep. I was feeling better minute by minute. My own blood, stored in bottles, had been returned to my body, and was now coursing through my arteries, my veins, reanimating me. We were going to make it.

Hell Notes (57 s) / Mr. Penumbra's Twenty-Four-Hour Book Store (65 s) (Incongruent)

It was the middle of a long, miserable winter. Icy arctic blasts alternated with stretches of unseasonable warmness when the snowy streets turn to rivers of gray mush. I wasted the morning on a couple of clients - an elderly pair who own two dozen down-home drive-ins dotted across rural Alabama. He wore a faded Hornets sweatshirt. She wore a pantsuit of polyester double-knit. They were on the verge of bankruptcy. They've been serving burgers and fries since the 50s. But now they were being crowded out by the big national chains. My agency was going to repackage them, give them an edge, bring them into the 21st century. With great gravitas, I swore to mercilessly tax every fiber of my creativity and marketing savvy until everything was coming up roses for them. Lying makes me hungry, so after that meeting I headed over to O'Reilly's for lunch.

I guess under general circumstances, this would feel like a creepy job requirement. Under the actual circumstances, selling rare books to mad scholars in the middle of the night, it feels perfectly appropriate. So, rather than spend my time staring at the forbidden shelves, I spend it writing about the customers. The basics, which book was purchased, its price, the time, go into the database. The rest goes into a giant, leather bound logbook. It's all mine. Mr. Penumbra pulled it out from under the front desk on my first night, heaved it open, and on the first page, he wrote my name. I have to say, I feel pretty proprietary about this book. I try to take clear, accurate notes, with only an occasional literary flourish. On quiet nights, I describe the weather. Sometimes I draw pictures, like Mr. Tindall tonight. So I guess you could say Rule Number Two isn't quite absolute. There's one book I'm allowed to touch in Penumbra's twenty-four-hour bookstore. It's the one I'm writing.

Love and Death in the Time of Monsters (60 s) / Blood of Virgins (58 s) (Incongruent)

Next to the TV were Mom's lighter and a pack of Marlboro Reds. Janie had been telling me for years I should make her quit. How could I do that? Some things you just can't tell your mom. Like the fact that she can't sing, or that margarine isn't really better for you than butter. Once, I watched her finish one cigarette, but she didn't have her lighter, so I was relieved when she was done. Then she took the dying cigarette and used it to light a fresh one. It was a clever use of fire, but I was horrified. I was six. I decided then I'd never smoke, years before our teachers lectured us on cancer. But the notebooks I brought to school smelled of tobacco. The summer after college, I couldn't find a job, and had to move back home. One night she asked me to run to the store and get her cigarettes. I didn't know what to do. They were poison, but she said she didn't want to live alone, now that Dad was gone. She wanted to be with him. She smoked because of love.

"I've got better things to do than spend an hour listening to some hippie chick complain that the world's not fair." "Aren't you curious what they'll say?"
"You can tell me all about it when you get back." The bathroom door closed behind him. So I went alone, and slipped into a chair at the back at the small auditorium. Out of 60 seats, only a dozen were taken. I guessed that the knot of five students in the first row were the campus Greens. Beside the podium hung a movie screen. Finally, at 8:15, when it was clear no stragglers would arrive, one of the Greens stood and faced the crowd. Matt would've snickered then, I was sure, would've whispered "I called it" if he'd bothered to come. The girl's mildly frizzy brown hair fell past her waist. She wore heavy black shoes, glasses with thick oval frames, and a homemade skirt of stitched-together red and blue and purple. A hippie chick. Matt would've laughed, but in her own way, she was beautiful.

Travels With My Cats (59 s) / End Game (62 s) (Incongruent)

That night I was faced with a major decision. I didn't want to read a book called "Travels with my Cats" by a woman called "Miss", but I'd spent my last nickel on it—well, the last until my allowance came due again next week—and I'd read all my other books so often you could almost see the eye tracks all over them. So, I picked it up without much enthusiasm and read the first page, and then the next, and suddenly I was transported to Kenya Colony, and Siam, and the Amazon. Miss Priscilla Wallace had a way of describing things that made me wish I was there, and when I finished a section, I felt like I'd been there. There were cities I'd never heard of before, cities with exotic names like Maracaibo and Samarkand and Addis Ababa. Some with names like Constantinople that I couldn't even find on the map. Her father had been an explorer. Back in the days when there still were explorers. She had taken her first few trips abroad with him, and he had undoubtedly given her the taste for distant lands.

Alan and I weren't friends, even though we sat across the aisle from each other—Edwards, Farr, Fitzgerald, Gallagher—and after the screaming fit, I thought he was just as weird as everyone else thought. I never taunted Alan like some of the boys, or laughed at him like the girls, and a part of me was actually interested in the strange things he sometimes said in class, always looking as if he had no idea how peculiar he sounded. But I wasn't strong enough to go against the herd, and make friends with such a loser. The summer before Alan went off to Harvard, we did become, if not friends, then chess companions. "You play rotten, Jeff," Alan said to me with his characteristic oblivious candor, "but nobody else plays at all." So, two or three times a week we sat on his parents' screened porch and battled it out on the chessboard. I never won. Time after time I slammed out of the house in frustration and shame, vowing never to return. After all, unlike wimpy Alan, I had better things to do with my time.

#### 4. WMS transcript

Anna [A] Thompson [B] of South [C] Boston [D] employed [E] as a cook [F] in a school [G] cafeteria [H] reported [I] at the police [J] station [K] that she had been held up [L] on State Street [M] the night before [N] and robbed [O] of fifty-six dollars. [P] She had four [Q] small children, [R] the rent was due, [S] and they had not eaten [T] for two days. [U] The police [V] touched by the woman’s story [W] took up a collection [X] for her [Y].

#### 5. *Pieman* Intact Story transcript

I began my illustrious career in journalism in the Bronx where I toiled as a hardboiled reporter for The Ram, the student Newspaper at Fordham University. And one day I’m walking to the campus center and out comes the elusive Dean McGowan, architect of a policy to replace Fordham’s traditionally working- to middle-class students with wealthier, more prestigious ones. So I whip out my notebook and I go up to him and I say "Dean McGowan, is it true that Fordham University plans to raise tuition substantially above the inflation rate? And if so, wouldn’t that be a betrayal of its mission?" And he stops and looks at me, and he says, "Listen up punk" And right then there’s a blur in the corner of my eye, which becomes this figure holding a cream pie, which becomes the guy standing next to me mashing a cream pie into Dean McGowan’s face. And then runs away. And the Dean is covered with cream. So I give him a moment. And I say "Dean McGowan, would you care to comment on this latest attack?" And the Dean says "Yes, I would care to comment... Fuck You!" So I race back to the newsroom with my scoop and I, and I find the editor, Jim Dwyer, who’s a senior—and he will go on to win a Pulitzer Prize, that is true. And he’s a big guy, I pitch him my story, I tell him what I’ve seen, and he says "Dean McGowan, that guy’s a dick, write it up." So I’m banging out my story and I know it’s good, and then I start to make it better by adding an element of embellishment. Reporters call this "making shit up". And they recommend against crossing that line. But I had just seen the line crossed between a high-powered Dean and assault with a pastry, and I kind of liked it. So the first thing I did was I gave the figure a name. I called him Pieman, capital P, capital M, and I described him as a cape-wearing masked avenger. Though in fact he’d been capeless. And I said that as he fled the scene, he clicked his heels in rakish glee. And I gave him a catch phrase, in Latin. I said that he cried out, "ego sum non un bestia", which means "I am not an animal", which makes no sense. I needed something... I’m Catholic ... Latin just comes to me. So I finish my story, and I hand it to Dwyer, and he reads it and he says, "Pieman, I love it, page one!" And that’s how the first line got crossed. Few days later I get a letter. It says, "Dear Jim, good story, nice details. If you want to see me again in action, be on the steps of Duane Library, Tuesday, at three o’clock. Signed, Pieman." So I was there with a photographer from The Ram. And sure enough, out comes Sheila Biel, student body president. And now Sheila was different from me and all of the other Fordham students who wore flannel shirts and worked part-time jobs. Sheila was well-bred. Sheila had school spirit. Sheila was the kind of student that Dean McGowan wanted more of. Although rumor had it that he got plenty of her in his office, on his desk, but that’s just a rumor. Please do not spread that outside this room. I myself would never say that. But the fact was, Sheila had collaborated with the Dean to ban outdoor drinking on campus. The infamous ‘No More Beer at Barbecues' rule - that’s right, boo that rule. Sheila thought drinking in public was in poor taste. I think you know what happened next. Pieman emerged from behind a late-night library drop box, made his delivery, and fled away, crying "ego sum non un bestia". Or at least that’s what it said in my story in the newspaper the next day, which ran with a photo of him leaving the scene, cape flowing behind him, doing this *clicks heels*... and that’s what made him a sensation on campus. People started dressing like him and quoting him in class. The Ram ran five major stories about Pieman, all of them by me. And towards the end of this run I was out at a bar one night and I saw, I came in and I saw in the corner, um, Angela, from my Brit Lit class, drinking with some friends. And now Angela and I had been flirting for two months. Or, I had been flirting with her. And with such nuance that there was a question about whether she knew I existed. So I saw her there and made a mental note to do nothing about it. And then I went to the bar to buy a round and I felt a tap on my shoulder. And it was her, and she said "Jim, we were just talking about how you always seem to know when and where Pieman will strike. And we were wondering, are you Pieman?" And I knew by the way she said it, I knew, that if I said yes, she would have sex with me. And wasn’t I really Pieman? For having brought him into being? Didn’t she only know about him through me? But she had asked me a straightforward question that came with a straightforward answer. In fact, I wasn’t Pieman. As far as I knew I had never seen the guy out of costume. So I looked at her and I said, "Yes, Angela, I am Pieman." And she said "Oh good, now buy me a beer and tell me all about it."

#### 6. *Pieman* Scrambled Paragraphs transcript

*Each paragraph’s order of appearance in the Intact Story: 5, 12, 3, 7, 6, 8, 11, 4, 10, 9, 2, 1*

5: who’s a senior—and he will go on to win a Pulitzer Prize, that is true. And he’s a big guy, I pitch him my story, I tell him what I’ve seen, and he says "Dean McGowan, that guy’s a dick, write it up." So I’m banging out my story and I know it’s good, and then I start to make it better by adding an element of embellishment. Reporters call this "making shit up".
[Duration: 30 seconds]

12: Or, I had been flirting with her. And with such nuance that there was a question about whether she knew I existed. So I saw her there and made a mental note to do nothing about it. And then I went to the bar to buy a round and I felt a tap on my shoulder. And it was her, and she said "Jim, we were just talking about how you always seem to know when and where Pieman will strike. And we were wondering, are you Pieman?" And I knew by the way she said it, I knew, that if I said yes, she would have sex with me. And wasn’t I really Pieman? For having brought him into being? Didn’t she only know about him through me? But she had asked me a straightforward question that came with a straightforward answer. In fact, I wasn’t Pieman. As far as I knew I had never seen the guy out of costume. So I looked at her and I said,
[Duration: 74 seconds]

3: standing next to me mashing a cream pie into Dean McGowan’s face. And then runs away. And the Dean is covered with cream. So I give him a moment.
[Duration: 17 seconds]

7: which means "I am not an animal", which makes no sense. I needed something... I'm Catholic ... Latin just comes to me.
[Duration: 13 seconds]

6: And they recommend against crossing that line. But I had just seen the line crossed between a high-powered Dean and assault with a pastry, and I kind of liked it. So the first thing I did was I gave the figure a name. I called him Pieman, capital P, capital M, and I described him as a cape-wearing masked avenger. Though in fact he'd been capeless. And I said that as he fled the scene, he clicked his heels in rakish glee. And I gave him a catch phrase, in Latin. I said that he cried out, "ego sum non un bestia",
[Duration: 47 seconds]

8: So I finish my story, and I hand it to Dwyer, and he reads it and he says, "Pieman, I love it, page one!" And that's how the first line got crossed. Few days later I get a letter. It says, "Dear Jim, good story,
[Duration: 20 seconds]

11: to ban outdoor drinking on campus. The infamous 'No More Beer at Barbecues' rule - that's right, boo that rule. Sheila thought drinking in public was in poor taste. I think you know what happened next. Pieman emerged from behind a late-night library drop box, made his delivery, and fled away, crying "ego sum non un bestia". Or at least that's what it said in my story in the newspaper the next day, which ran with a photo of him leaving the scene, cape flowing behind him, doing this *clicks heels*... and that's what made him a sensation on campus. People started dressing like him and quoting him in class. The Ram ran five major stories about Pieman, all of them by me. And towards the end of this run I was out at a bar one night and I saw, I came in and I saw in the corner, um, Angela, from my Brit Lit class, drinking with some friends. And now Angela and I had been flirting for two months.
[Duration: 72 seconds]

4: And I say "Dean McGowan, would you care to comment on this latest attack?" And the Dean says "Yes, I would care to comment... Fuck You!" So I race back to the newsroom with my scoop and I, and I find the editor, Jim Dwyer,
[Duration: 20 seconds]

10: the other Fordham students who wore flannel shirts and worked part-time jobs. Sheila was well-bred. Sheila had school spirit. Sheila was the kind of student that Dean McGowan wanted more of. Although rumor had it that he got plenty of her in his office, on his desk, but that's just a rumor. Please do not spread that outside this room. I myself would never say that. But the fact was, Sheila had collaborated with the Dean
[Duration: 31 seconds]

9: nice details. If you want to see me again in action, be on the steps of Duane Library, Tuesday, at three o'clock. Signed, Pieman." So I was there with a photographer from The Ram. And sure enough, out comes Sheila Biel, student body president. And now Sheila was different from me and all
[Duration: 26 seconds]

2: is it true that Fordham University plans to raise tuition substantially above the inflation rate? And if so, wouldn't that be a betrayal of its mission?" And he stops and looks at me, and he says, "Listen up punk" And right then there's a blur in the corner of my eye, which becomes this figure holding a cream pie, which becomes the guy
[Duration: 23 seconds]

1: [end of music] I began my illustrious career in journalism in the Bronx where I toiled as a hardboiled reporter for The Ram, the student Newspaper at Fordham University. And one day I'm walking to the campus center and out comes the elusive Dean McGowan, architect of a policy to replace Fordham's traditionally working- to middle-class students with wealthier, more prestigious ones. So I whip out my notebook and I go up to him and I say "Dean McGowan,
[Duration: 33 seconds]

*13: "Yes, Angela, I am Pieman." And she said "Oh good, now buy me a beer and tell me all about it."
[Duration: 18 seconds; discarded in the scrambled stimulus due to short duration]

**Supplementary Note 2: D. A.'s post *Pieman* scan interview transcript**

Experimenter: So how did you find it, in the—

D. A.: Yeah, a little unusual, it's kinda a bit strange but then it's, uh, uh, I think it's interesting, this brain, to find the patterns and images other than just—

Experimenter: Absolutely.

D. A.: —rather than just listening to your talk or voices all in.

Experimenter: Exactly, yeah, yeah, absolutely. So well because they're memories—

D. A.: Changes your focal point.

Experimenter: Absolutely.

D. A.: Where everything is.

Experimenter: Absolutely. Because it emerges from the brain, so we were able to see it, and you kept so still, with the pictures we got were so perfect, so thank you for—

D. A.: It was, it was, it was an inducement to stay still and to pay attention [laughs].

Experimenter: Yeah, you did stay still well. Only if all our participants could be, could be, so good. So, I'm just gonna ask you a few quick questions about when you were in the scanner, there was a story, um do you, you listened to a story. Do you remember anything about it? Can you tell me—

D. A.: Details um might have been about a young girl or boy trying to go, wanting to go out into the city and, and to work um.

Experimenter: Cool, yup.

D. A.: And, uh.

Experimenter: Great.

D. A.: Yeah, that's. Details are getting…

Experimenter: Great. Ok. So I can give, I'm just gonna give you a few, you know, hints. So it was set on a university campus.

D. A.: A campus.

Experimenter: Do you, yeah, does that ring a bell or?

D. A.: Uh, hmm, should it be in resident or be in, kind of, live off campus and …

Experimenter: Yeah, yeah.

D. A.: …work or…

Experimenter: Yeah, yeah, great. And there was a college newspaper—

D. A.: Hmm.

Experimenter: —in the story. Does that—?

D. A.: And they wanted to interview him and, and get opinions on him, this, what the campus is like and what it is to go, is it a good campus and a good university, yeah.

Experimenter: Great, great. Um, and um if I told you that there was a character in there called Pieman, does that bring—

D. A.: Hmm.

Experimenter: —does that bring that to anything?

D. A.: It, it brings something up in my mi–mind, but I'm not sure if he was kind of like the local town clown or the bad campus clown [laughs].

Experimenter: Ok, cool, great, great, cool. And then the last, there was a dean at the college who was a character as well. Does that—?

D. A.: Uh, no.

Experimenter: No. Ok, excellent. Great.

D. A.: Unless it's Dean Weatherby [laughs].
